# Phage-Assisted, Active Site-Directed Ligand Evolution of a Potent and Selective Histone Deacetylase 8 Inhibitor

**DOI:** 10.1101/2022.09.14.508049

**Authors:** Jared S. Morse, Yan J. Sheng, J. Trae Hampton, Lauralee D. Sylvain, Sukant Das, Yugendar R. Alugubelli, Peng-Hsun Chase Chen, Kai S. Yang, Shiqing Xu, Carol A. Fierke, Wenshe Ray Liu

## Abstract

The phage-assisted, active site-directed ligand evolution (PADLE) is a recently developed technique that uses an amber codon-encoded noncanonical amino acid (ncAA) as an anchor to direct phage-displayed peptides to a target for an enhanced ligand identification process. 2-Amino-8-oxodecanoic acid (Aoda) is a ketone-containing ncAA residue in the macrocyclic peptide natural product apicidin that is a pan-inhibitor of Zn^2+^-dependent histone deacetylases (HDACs). Its ketone serves as an anchoring point to chelate the catalytic zinc ion in HDACs. Using a previously evolved *N*^*ε*^*-*acetyl-lysyl-tRNA synthetase in combination with tRNA^Pyl^, we showed that Aoda was efficiently incorporated into proteins in *Escherichia coli* by amber suppression. By propagating an amber codon-obligate phagemid library in *E. coli* encoding Aoda, we generated an Aoda-containing phage-displayed peptide library. Using this library to conduct PADLE against HDAC8 revealed a 7-mer peptide GH8P01F1 with Aoda-flanking amino acid residues that match existing peptide sequences in identified HDAC8 substrates. Switching Aoda in GH8P01F1 to a more Zn^2+^-chelating ncAA *S*-2-amino-8-hydroxyamino-8-oxooctanoic acid (Asuha) led to an extremely potent GH8HA01 that has an HDAC8-inhibition K_i_ value as 0.67 nM. GH8HA01 and its 5-mer truncation analogue Ac-GH8HA01*Δ*1*Δ*7 that has an HDAC8-inhibition K_i_ value as 0.31 nM represent two most potent HDAC8 inhibitors that have been developed so far and both are highly selective against HDAC8 over the other three tested HDACs, demonstrating the great potential of using PADLE to identify highly potent and selective ligands for targets.

## 1. INTRODUCTION

Histone deacetylases (HDACs) are epigenetic erasers which play a pivotal role in regulating the structure and function of histones and other proteins. Aberrant HDAC activity is associated with various cancers, neurodegenerative diseases, and many other disease states.^1-3^ These enzymes are grouped into four classes: three Zn^2+^-dependent classes (I, II, and IV) and one NAD^+^-dependent class (III, also called Sirtuins). The Zn^2+^-dependent classes are grouped based off their homology to corresponding yeast protein deacetylases and share a rather conserved active site structure. Many of these HDACs have become highly desirable drug targets for their therapeutic potential in treating cancers, infection, neurodegeneration and more.^4^

The traditional paradigm of Zn^2+^-dependent HDAC inhibition utilizes three motifs: a zinc binding group (ZBG) that positions the inhibitor in the active site, a linker that binds within the active site channel, and a capping group that forms interactions at the surface of the HDAC enzyme. Despite the distinct substrate profile of the different HDACs, their sequences are highly conserved, especially within the active site tunnel.^5^ Much of the rational drug design has centered around enhancement of the ZBG, starting with the chelating hydroxamate in vorinostat and further optimization for selectivity or safety.^6,7^ Many HDAC inhibitors have been reported, including vorinostat and belinostat, which are pan-inhibitors that do not discern between the individual isoforms of HDACs.^8^ Other reported inhibitors, including the FDA approved romidepsin and panobinostat, display selectivity against one or two classes of HDACs. The conserved nature of the HDAC active site channel, which natively recognizes the *N*^*ε*^-acetyl-L-lysine sidechain of protein substrates, limits the ability to exploit this region for isoform selectivity in the design of HDAC inhibitors. Additionally, the enzyme surface adjacent to the active site is malleable and accommodates ligands in a different, context-dependent manner based upon the identity of the binding partner.^9,10^ These two facts combined result in the observed difficulty in designing HDAC inhibitors which display selectivity at the isoform level. Interactions distal to the active site channel are important for substrate specificity and are exploitable for inhibition. For example, some cyclic peptides without a ZBG can still act as competitive inhibitors while retaining class I selectivity.^11^ Therefore, methods for screening novel capping groups can greatly aid in the development of selective HDAC inhibitors.

The class I HDAC8 has long been associated with various cancer subtypes, but only recently has there been elucidation in the mechanism behind HDAC8-associated disease progression.^12-14^ HDAC8’s mechanistic possibilities extend beyond simply regulating gene expression, as it has been found to localize in the cytoplasm as well as the nucleus and evidence grows for the existence of non-histone substrates, such as p53.^15-17^ As evidence grows for HDAC8’s role in disease progression, so too grows the desire for potent and selective HDAC8 inhibitors. The pan-HDAC inhibitor quisinostat (HDAC IC_50_s: 0.11 to 119 nM; HDAC8 IC_50_: 4.26 nM) potently inhibits HDAC8, but failed to demonstrate superior results compared to vorinostat or romidepsin in phase 2 clinical trials for treating T-cell lymphoma.^18-20^ PCI-34051 is a highly selective HDAC8 inhibitor with a K_i_ = 10 nM.^12^ Its selectivity is achieved by taking advantage of a unique feature in the active site pocket of HDAC8 with flexible L-1 and L-6 loops that are otherwise sterically hindersome in other class I HDACs.^21^ So far, there are no inhibitors that achieve selectivity by interactions at the surface of HDAC8.

Our group has previously published on a newly-developed phage-assisted, active site-directed ligand evolution (PADLE) technique that utilizes an amber codon-obligate library to genetically encode a non-canonical amino acid (ncAA) as a warhead into phage-displayed peptides to direct them towards the active site of a target enzyme or protein.^22^ In contrast to traditional phage display methods, which are agnostic to target protein-ligand interactions, PADLE increases the rate of discovery of productive peptide binders by stacking the deck towards competitive binding guided by the ncAA ligand. The amber codon-obligate library eliminates the inherent bias of peptides with and without ncAA in phage production and propagation, maximizing the effect of target binding during the selection process, in contrast to previous ncAA-displaying phage peptides which necessitated strong ncAA-target binding to overcome this bias.^23^ In this paper, we use PADLE in combination with an ncAA warhead that acts as an *N*^*ε*^-acetyl-L-lysine isostere to guide peptide selection towards the active site tunnel, while the flanking randomized peptides are able to take advantage of the same interactions upon which substrate specificity arises, to evolve a potent and selective HDAC8 inhibitor. Following the generation of a singly enriched peptide sequence, subsequent structure activity relationship (SAR) studies and optimization of the ZBG resulted in the first sub-nanomolar selective HDAC8 inhibitor.

## 2. RESULTS AND DISCUSSION

### 2.1 The genetic incorporation of 2-amino-8-oxodecanoic acid (Aoda)

We selected HDAC8 for PADLE-based ligand optimization due to its implication in various cancers.^24-27^ For PADLE-based ligand search of HDAC8 inhibitors, an initial ligand that binds to the HDAC8 active site is necessary. Although an ncAA with a hydroxamate side chain could be potentially used as a starting ligand, we decided not to pursue this route due to the demonstrated super potency of hydroxamate to chelate the Zn^2+^ ion in the catalytic center that potentially obscures other interactions. A ligand with relatively weak binding is a better choice. For this reason, we set our sight on the naturally occurring cyclic tetrapeptide apicidin (Figure 1a). Apicidin is a pan-HDAC inhibitor that inhibits HDAC8 with a determined IC_50_ value of 2.53 µM.^28^ It contains Aoda, a facile ZBG that acts as an *N*^*ε*^-acetyl-L-lysine isostere. As an ncAA, Aoda (Figure 1b) is a weak electrophile in solution, but is hydrated in the active site of HDAC8 and forms a geminal diol that coordinates to the Zn^2+^ ion in the active site.^29^ Aliphatic ketones rarely generate their hydrated forms in solution (K_eq_ < 0.002 for hydration of acetone),^30^ and therefore the ZBG is only “active” when it is positioned within the HDAC8 active site. We reasoned that a genetically incorporated Aoda that can serve as a weak ligand for HDAC8 in a phage-displayed peptide library will direct coded peptides to the active site of HDAC8 for the selection of more potent and highly selective HDAC8-targeting peptides since residues flanking on the two sides of Aoda can potentially engage HDAC8 around but not at the active site to generate selective interactions at the enzyme surface. In this manner, the peptide library can be selected to exploit the same interactions upon which substrate specificity arises.

**Figure 1:**
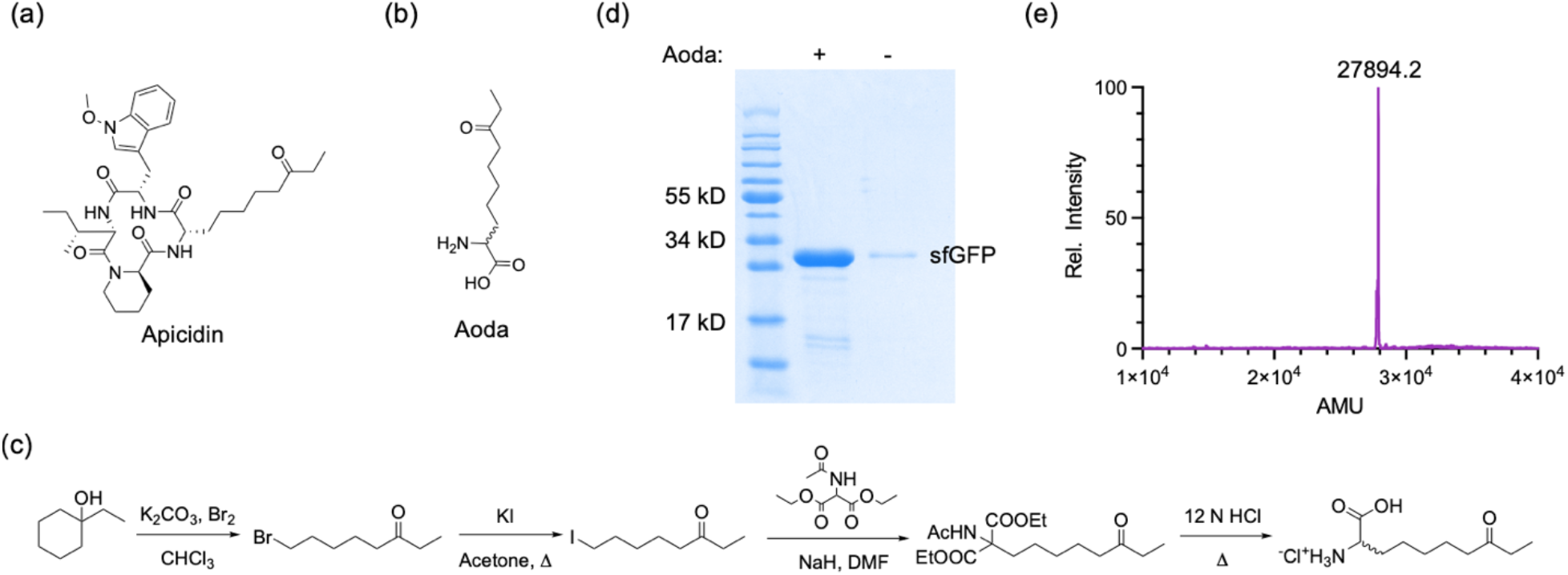
a) Structure of the macrocyclic peptide apicidin, a pan-HDAC inhibitor. b) Structure of the ncAA Aoda. c) Synthetic route for the hydrochloride salt of Aoda. d) SDS-PAGE of sfGFP-134TAG expressed in the presence/absence of 4 mM Aoda. Cells were transformed with two plasmids encoding MmAcKRS, tRNA^Pyl^, and sfGFP134TAG. e) ESI-MS of sfGFP with Aoda in the 134^th^ amino acid position. Calculated mass: 27,894.9 AMU.

To demonstrate this prospect, we synthesized Aoda as the racemate (Figure 1c) and tested its genetic encoding by amber codon. Although Aoda was a known compound with established synthetic routes, we designed a much simplified synthetic procedure that involved a retro-Barbier fragmentation reaction to directly convert 1-ethylcyclohexan-1-ol to 8-bromooctan-3-one. 8-Bromooctan-3-one was converted to an iodide form and then reacted with diethyl 2-acetamidomalonate followed by hydrolysis to generate Aoda. With this new synthetic route, we were able to achieve the synthesis of Aoda with the overall yield as 52% in just four steps. We decided to use the racemate instead of the chiral L-form of Aoda for our following cellular demonstration of Aoda incorporation due to the preferential recognition of L-amino acids by the ribosome. In order to genetically incorporate Aoda by amber suppression, we used a previously evolved *M. mazei* pyrrolysyl-tRNA synthetase (MmAcKRS) and its cognate tRNA^Pyl^.^31^ The MmAcKRS-tRNA^Pyl^ was demonstrated to incorporate other ncAAs including *N*^*ε*^-acryloyl-L-lysine and *N*^*ε*^-propionyl-L-lysine as well.^32,33^ Due to the similarity between *N*^*ε*^-propionyl-L-lysine and Aoda, we suspected that MmAcKRS would also recognize Aoda as a substrate. To demonstrate this, we transformed *E. coli* BL21(DE3) cells with two plasmids, pEVOL-MmAcKRS that encoded MmAcKRS and tRNA^Pyl^ and pBAD-sfGFP134TAG that encoded superfolder green fluorescent protein (sfGFP) with an amber mutation at the 134^th^ amino acid coding position. Transformed cells were grown in the presence or absence of 4 mM Aoda. If MmAcKRS is able to charge tRNA^Pyl^ with Aoda, it can mediate the suppression of the amber mutation at the 134^th^ position of the sfGFP gene leading to the expression of full-length sfGFP. Otherwise, a truncated protein will be expressed. As expected and shown in Figure 1d, full-length sfGFP (sfGFP134Aoda) was only expressed in the presence of 4 mM. The expression yield was 34 mg/L in the 2YT medium. This expression level is comparable to that for other ncAAs such as *N*^*ε*^-Boc-L-lysine mediated by the wild-type PylRS.^34^ In the absence of Aoda, full-length sfGFP was expressed at a minimal level that matched what we typically observed as background suppression of amber codon when the MmAcKRS-tRNA^Pyl^ pair was used.^31^ We purified sfGFP134Aoda and characterized it using electrospray ionization mass spectrometry (ESI-MS). Its deconvoluted spectrum showed an average molecular weight as 27,894.2 Da that matched well with its calculated average molecular weight as 27,894.9 Da (Figure 1E). This represented Aoda in its unhydrated form. Since MmAcKRS is able to mediate efficient incorporation of Aoda, we did not optimize the enzyme further for the Aoda incorporation and used it directly to build an Aoda-incorporated phage display library.

### 2.2 The construction of an Aoda-containing phage display library and its selection against HDAC8

After successful demonstration of the genetic incorporation of Aoda by amber codon, we proceeded to use the system to construct an Aoda-containing phage display library with randomized 7-mer peptides. M13 phages were expressed using TOP10 cells which were transformed with the following 3 plasmids: (1) pADL(NNK)_7_gIII, which encoded a 7-mer amber codon-obligate peptide library at the N-terminus of the pIII protein,^22^ (2) pEDF-AcKRS,^33^ a previously constructed plasmid that contains the genes encoding MmAcKRS and tRNA^Pyl^ for the site-specific incorporation of Aoda at amber codon within the 7-mer library, and (3) M13KO7-gIIITAA,^33^ a helper phage plasmid with a deleterious mutation at gIII and a modified M13 origin of replication which aids in the preferential packaging of the pADL(NNK)_7_gIII phagemid into phage particles whose displayed peptide phenotypes matched their packaged phagemid DNA genotypes.^35^ The amber-obligate pIII library was previously prepared via an invented superinfection-immunity selection approach that enriched a randomized 7-mer peptide-coding library to contain at least one amber codon for coding a genetically encoded ncAA.^22^ The use of this amber codon-obligate library was to avoid the growth disadvantage of amber codon-containing clones over non-amber codon clones. Transformed Top10 cells were grown in the 2YT medium with 10 mM Aoda to produce phages. 10 mM Aoda was used to boost up the phage production. Produced Aoda-containing phages were then used to select against GST-HDAC8 that was expressed in insect cells.

GST-HDAC8 was immobilized on glutathione-coating magnetic beads and used for biopanning against the Aoda-presenting phages. Iterative rounds of selection were conducted by introducing increasingly more stringent selection pressures (Figure 2). In the second round a negative selection against glutathione S-transferase (GST) immobilized on the same beads was introduced, and in the third round an extra wash step was added. The effective library size (3.88 × 10^9^ transformants of the amber codon-obligate (NNK_7_) library) and the initial phage titer for the first round of selection (7 × 10^9^ cfu) were on the same order of magnitude. Thus, it’s expected that the most probable number of any individual peptide sequence would be 1-2 copies. For this reason, the selection pressure was minimized in the first round to prevent the loss of productive binding sequences through statistical chance (e.g., unbound during a washing step, retained during a transfer step, etc.). Gradually increasing the selection pressure allowed for greater selection pressures to separate strong and weak binding sequences after the library was enriched to such an extent that each experiment is expected to contain multitudes more copies of a given sequence, where such statistical chances are negligible to the enrichment of a given peptide sequence. Phage production in the first expression increased 23-fold in the presence of 10 mM Aoda versus phages expressed in the absence of ncAA, indicating successful incorporation of Aoda into pIII (Figure S1). This ratio increased through the three rounds of selection against GST-HDAC8, where phages after the third round were produced 50-fold more in the presence of Aoda, providing indirect evidence of the further enrichment of the amber codon-containing clones over phages containing canonical sequences. After three rounds of selection, the enriched library was sequenced using Illumina next generation sequencing (Table 1). The selection converged to a single, highly enriched sequence: YXFFWED (X: Aoda) that was subsequently named as GH8P01. We observed that this sequence had structural similarities to previously discovered peptide substrates and inhibitors of HDAC8. Relative to the side chain positioned in the active site, this peptide shares a Phe in the +1 position and a Trp in the +3 position, consistent with our previous work which found the peptide Ac-FGXFSW-NH_2_ (X = *N*^*ε*^-acetyl-L-lysine), based on ZNF318, as one of the most efficient peptide substrates for HDAC8.^36^ Trapoxin A is a natural product HDAC inhibitor that bears an α,β-epoxyketone functional group as a ZBG. It contains also a Phe in the +2 position which forms end-on π-π interactions with a second Phe in the +1 position as well as with Y100 on HDAC8.^37^ Previous structural characterizations of HDAC8 bound to small molecule inhibitors have demonstrated that HDAC8 accommodates different conformations of small molecule inhibitors primarily mediated by the L1 loop which forms the surface area of one face of the active site channel and extends to the protein surface.^9^ Other regions, such as the L2 loop, can become more ordered when bound to certain ligands and remain disordered when bound to others. The L2 loop contains Y100 that forms π-π interactions with aromatic capping groups in both small molecules and macrocyclic peptides. It is likely that GH8P01 engages similar interactions with Y100 of HDAC8.

**Table 1:**
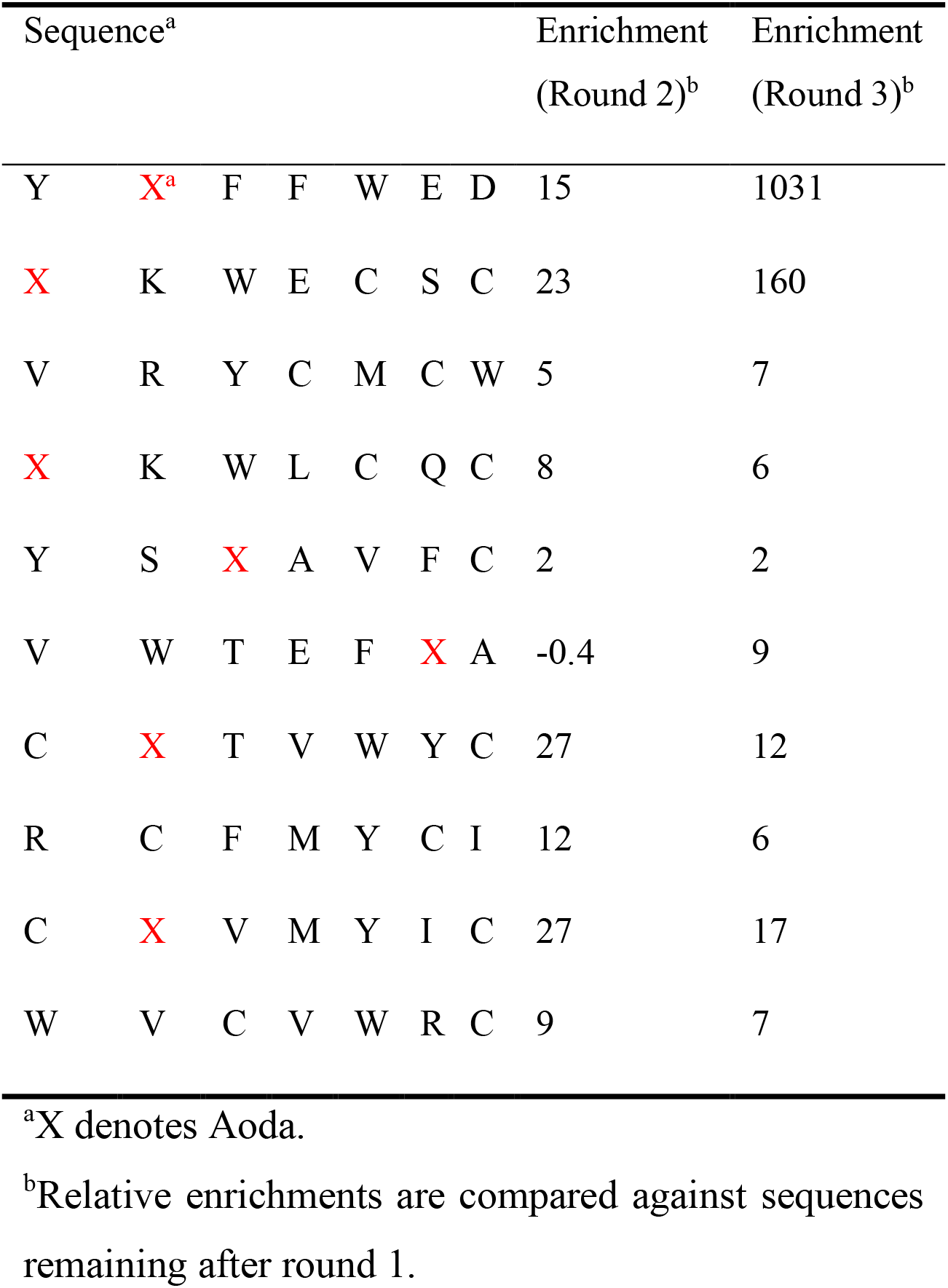
The most prevalent sequences after three rounds of selection against GST-HDAC8

**Figure 2:**
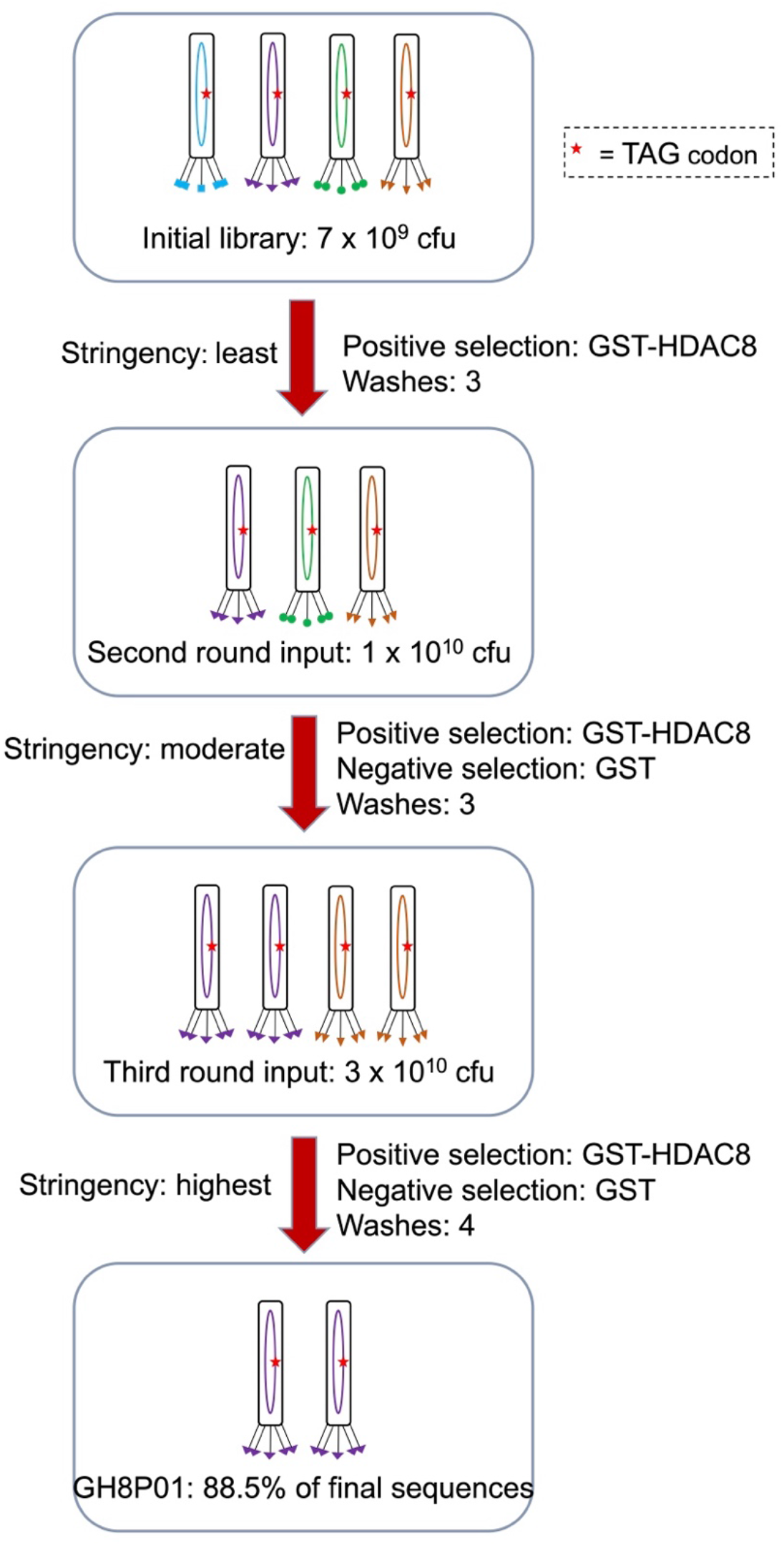
Selection strategy over the course of three rounds. An initial library displaying peptides containing Aoda were biopanned against immobilized GST-HDAC8 with minimized selection pressure to maximize retention of bound phages. The second round introduced a negative selection against immobilized GST to remove phages bound to the GST tag, and the third round introduced an additional wash. This process resulted in convergence to a single highly enriched peptide.

### 2.3 Validation of Aoda-containing enriched peptides as HDAC8 inhibitors

A 7-mer peptide based off GH8P01 was chemically synthesized using racemic Fmoc-Aoda. Fmoc-Aoda was synthesized from racemic Aoda by reacting it with Fmoc-OSu and then used in solid-phase peptide synthesis (SPPS). SPPS resulted in the creation of two major diastereomers that were separated via reverse phase HPLC and termed GH8P01F1 and GH8P01F2 based upon their order of elution. Previous reports of selections against GST-tagged proteins reported false positive ligands which bound to the GST-target fusion protein but not to tag-free target protein nor to GST enzyme by itself.^38^ To confirm peptide binding was not contingent upon the GST-HDAC8 fusion, all inhibition assays were performed with tag-free recombinant HDAC8 that was expressed in *E. coli* cells by following previous published protocols from us and others.^10,39,40^ HDAC8 was purified in a metal-free procedure as the apo enzyme and then reconstituted with one equivalent of Zn^2+^ to maximize its enzyme activity. Enzymatic activity was measured in an activity assay utilizing a fluorogenic peptide substrate Ac-LG-K(Tfa)-AMC which produces a signal upon addition of a developing solution if the substrate’s C-terminal *N*^*ε*^-trifluoroacetyl-L-lysine (K(Tfa)) is deacylated.^20,41^ Although Ac-LG-K(Tfa)-AMC was reported before, we optimized its synthesis by following a synthetic route presented in Scheme S3. Ac-LG-K(Tfa)-AMC was purified to more than 95% purity and then used in our kinetic analysis. In the kinetic analysis, we used 0.4 nM HDAC8, 50 *µ*M Ac-LG-K(Tfa)-AMC, and varied concentrations of GH8P01F1 or GH8P01F2 at pH 8. Based on the inhibition assay results, GH8P01F1 was found to be an active inhibitor of HDAC8 with an IC_50_ value of 5.3 ± 1.0 µM, while GH8P01F2 was a poor inhibitor showing only ∼40% inhibition of HDAC8 at 25 µM (Figure 3a). Since ribosomes recognize L-amino acids much more efficiently than D-amino acids, we assigned that GH8P01F1 contains the L-isomer of Aoda as the ZBG.

**Figure 3:**
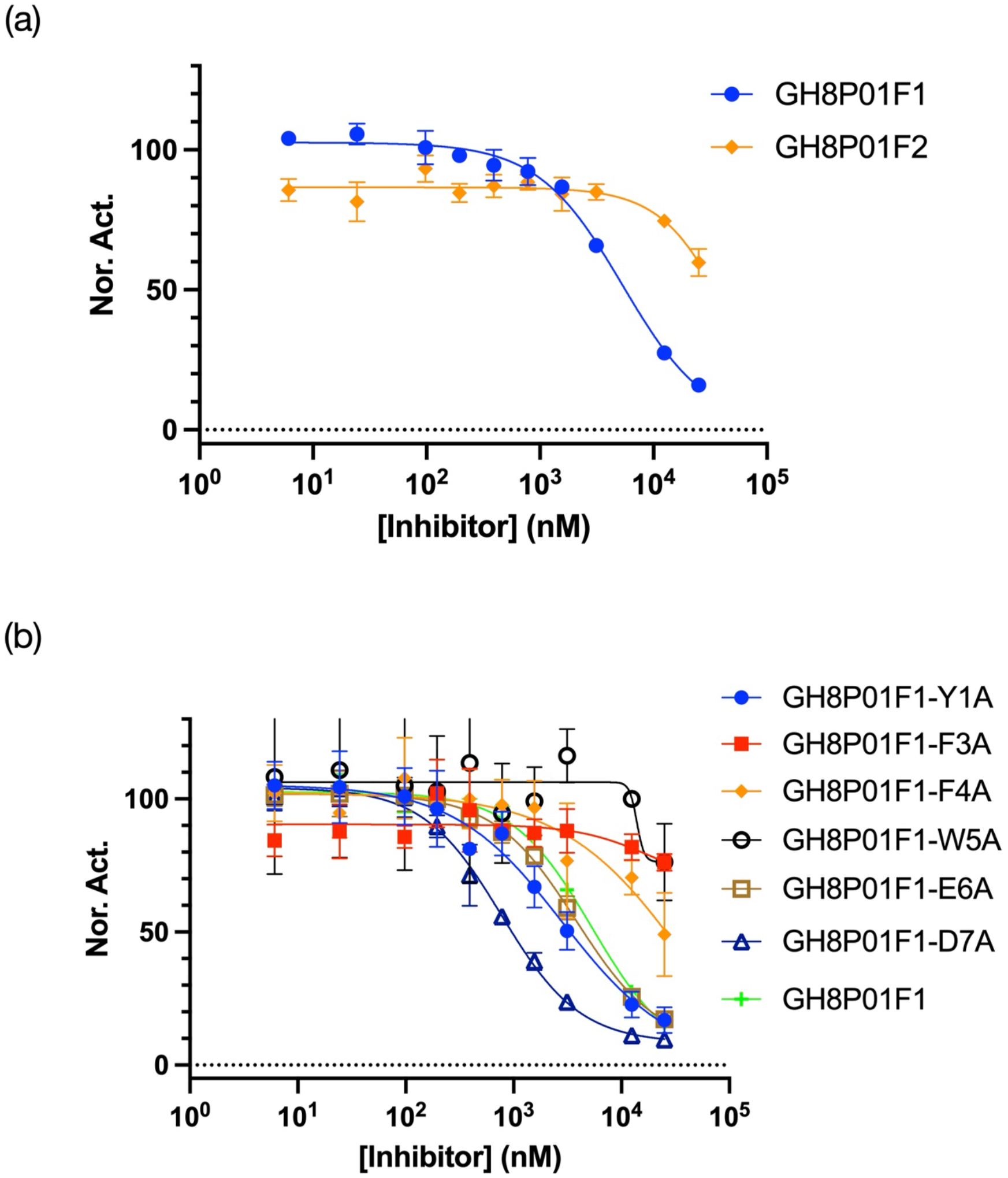
a) IC_50_ curves of the Aoda-containing HDAC8 inhibitors GH8P01F1 and GH8P01F2. IC_50_ curves of GH8P01F1 derivatives. An alanine scan was performed by iteratively replacing each canonical residue with an alanine and testing for change of HDAC8 inhibition.

We next wanted to optimize GH8P01 to be a more potent HDAC8 inhibitor and a better molecular probe. For this purpose, an alanine scan was performed (Figure 3b) to further study the enriched peptide sequence and the contribution that each residue makes towards binding to HDAC8. Replacing the 3^rd^, 4^th^, or 5^th^ residue of GH8P01 with an alanine resulted in a significant decrease in inhibition, indicating that these aromatic residues are vital for binding to and recognizing HDAC8 (summarized in Table 2). Replacing the 1^st^ or 6^th^ residues had a slight increase in inhibition (2.8 and 3.8 vs 5.3 µM IC_50_). Additionally, replacement of the 7^th^ residue gave a noticeable increase in inhibition (0.77 vs 5.3 µM IC_50_). Based on these results, it is our belief that some residues in GH8P01 were enriched not because they enhanced binding to the target HDAC8 protein, but rather aided in phage propagation. We have reported a previous selection done with the same 7-mer library.^22^ Its enriched peptide sequences could be truncated with minimal perturbation to inhibition. Notably, the 7^th^ code position in our TAG-obligate randomized library was disproportionately less-enriched for the TAG codon, which could indicate a role for this residue in phage propagation.^22^ For residues which do not directly contribute to phage-target interactions, “fitness” in the context of a selection will be determined by factors beyond binding affinity but still contribute to an individual phage’s propensity to replicate and propagate through the biopanning experiment.

**Table 2:** IC50 values of the enriched Aoda-containing peptides and alanine scan derivatives against HDAC8

| Peptide | Sequence |  |  |  |  |  |  |  | IC<sub>50</sub> (μM) |
| --- | --- | --- | --- | --- | --- | --- | --- | --- | --- |
| GH8P01F1 | Y | X | F | F | W | E | D |  | 5.3 ± 1.0 |
| GH8P01F2 | Y | X | F | F | W | E | D |  | 40%<sup>a</sup> |
| GH8P01F1-Y1A | A | X | F | F | W | E | D |  | 2.8 ± 1.2 |
| GH8P01F1-F3A | Y | X | A | F | W | E | D |  | 24%<sup>†</sup> |
| GH8P01F1-F4A | Y | X | F | A | W | E | D |  | 51%<sup>†</sup> |
| GH8P01F1-W5A | Y | X | F | F | A | E | D |  | 24%<sup>†</sup> |
| GH8P01F1-E6A | Y | X | F | F | W | A | D |  | 3.8 ± 0.6 |
| GH8P01F1-D7A | Y | X | F | F | W | E | A |  | 0.77 ± 0.07 |

<sup>a</sup>Inhibition reported at 25 μM of peptide

### 2.4 Replacement of the ZBG with *S*-2-amino-8-hydroxyamino-8-oxooctanoic acid (Asuha)

Since many HDAC inhibitors engage the active site Zn^2+^ ion with a strong chelating group, we wanted to explore whether replacing the Aoda residue in GH8P01 with a stronger ZBG warhead with similar length will lead to a more potent inhibitor. Asuha is an L-amino acid with a side chain hydroxamic acid that has been observed in many potent and clinically approved HDAC inhibitors. Since Asuha and Aoda contain identical aliphatic linkers, we think replacing Aoda in GH8P01 with Asuha will potentially generate a more potent HDAC8 inhibitor. In this newly designed peptide termed GH8HA01, the side chain keto group of Aoda is replaced by the hydroxamic acid in Asuha. To conduct SPPS of GH8HA01, we synthesized Fmoc-Asu(NHOPMB)-OH. Fmoc-Asu(NHOPMB)-OH was synthesized from Fmoc-Asu (Asu: 2-amino-suberic acid) by coupling with O-(4-methoxybenzyl)hydroxylamine. GH8HA01 was produced as a chiral pure compound with only an L-amino acid in the metal chelation site. As predicted, this peptide was drastically more potent than GH8P01 and inhibited HDAC8 with an IC_50_ of 0.85 ± 0.13 nM (Figure 4a). Following the insight gained by the alanine scan with Aoda-containing peptides, we synthesized three new analogues of GH8HA01 by removing the Asp residue (GH8HA01Δ7), replacing the 1^st^ Tyr residue with an acetyl capping group (Ac-GH8HA01Δ1), and a pentameric derivative with both modifications (Ac-GH8HA01Δ1Δ7). The first two derivatives displayed only a slight loss in inhibition compared to GH8HA01, reaffirming the roles of the residues in the 1^st^ and 7^th^ position as ineffectual at contributing to HDAC8 binding. The last peptide displayed remarkable inhibition with IC_50_ as 0.39 ± 0.05 nM, half that of the parent peptide. The IC_50_s of these hydroxamate peptides are summarized in Table 2. As far as we know, Ac-GH8HA01Δ1Δ7 is the most potent HDAC8 inhibitor that has been discovered.

**Figure 4:**
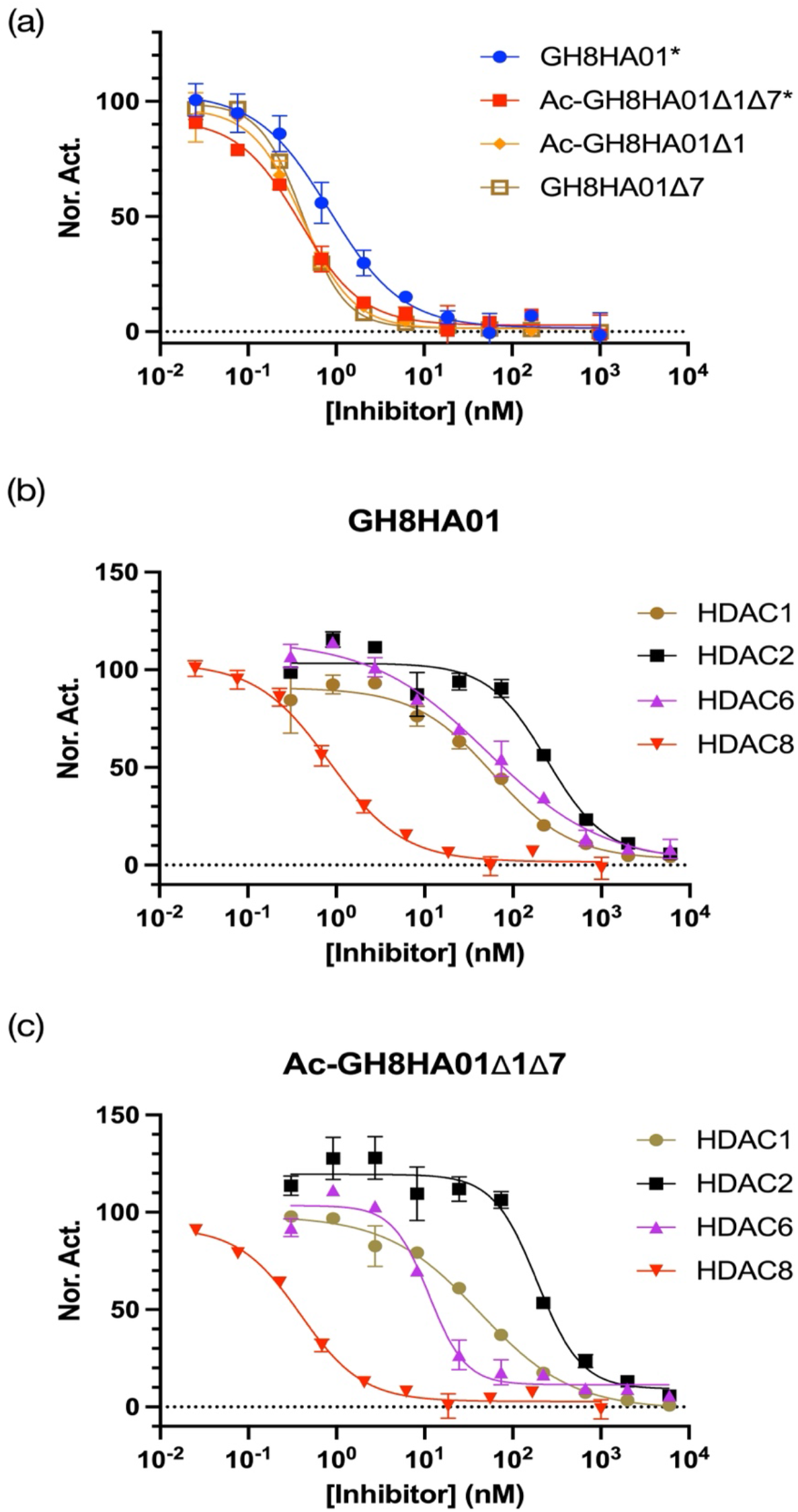
a) IC_50_ curves of the Asuha-containing HDAC8 inhibitor GH8HA01 and its truncated derivatives. b) IC_50_ curves of the 7-mer peptide inhibitor GH8HA01 against HDAC1, 2, 6, and 8. IC_50_ curves of the 5-mer peptide inhibitor Ac-GH8HA01Δ1Δ7 against HDAC1, 2, 6, and 8.

### 2.5 Selectivity of GH8HA01 and Ac-GH8HA01Δ1Δ7 toward HDAC8 over other HDAC enzymes

To confirm the selectivity of our peptides, further inhibition assays were performed with other HDAC isoforms including HDAC1, HDAC2 and HDAC6 (Figure 4). Enzymatic activity was measured using a different fluorogenic substrate, Ac-LG-K(Ac)-AMC. This previously reported substrate for HDACs 1, 2, and 6 was synthesized according to the route presented in Scheme S4. The kinetic analysis was done with various enzyme concentrations (HDAC1: 0.8 nM; HDAC2: 2.4 nM: HDAC6: 0.8 nM) in HEPES buffer at pH 8.0, with 20 µM Ac-LG-K(Ac)-AMC, and varied concentrations of peptide inhibitors. GH8P01F1 showed no detectable inhibition against all three HDACs tested at 25 *µ*M. GH8HA01 and Ac-GH8HA01Δ1Δ7 were tested for their inhibition against each HDAC and an IC_50_ value was measured for each peptide. The corresponding inhibition constants (K_i_) were estimated using the Cheng-Prusoff equation (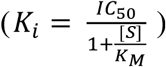) and previously reported Michaelis-Menten constants (HDAC1: K_M_ = 6 µM, HDAC2: K_M_ = 3 µM, HDAC6: K_M_ = 16 µM, HDAC8: K_M_ = 190 µM).^41,42^ These K_i_ values were used to determine the selectivity of each peptide for HDAC8 over other HDAC isoforms (Table 3). The hydroxamic acid-containing peptides both showed remarkable selectivity, with GH8HA01 displaying 19-fold selectivity against HDAC1 and higher selectivity against HDAC 2 and 6. Ac-GH8HA01Δ1Δ7 displayed 16-fold selectivity against HDAC6 and even higher selectivity against the class I HDACs tested. Therefore, the PADLE-based selection followed by the ZBG functional group conversion is a practical method to develop both potent and selective HDAC8 inhibitors.

**Table 3:**
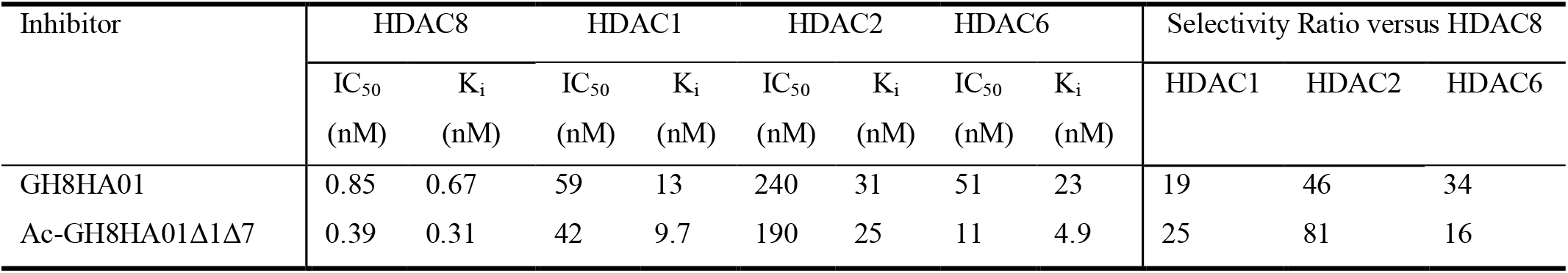
Inhibition constants and HDAC8 selectivity of hydroxamate-containing peptides against various HDAC isoforms

## 3. CONCLUSION

PADLE has provided a modular platform to genetically incorporate a known ligand into a peptide library and quickly enrich selective and potent peptide inhibitors. In our previous work, we demonstrated an application of PADLE in the identification of potent and selective SIRT2 inhibitors.^22^ In the current study, we used Aoda to guide displayed peptides towards the active site of HDAC8 and successfully identified the low-micromolar inhibitor GH8P01F1. With small optimizations to the sequence and ZBG, we report the discovery of two sub-nanomolar HDAC8 inhibitors: the heptamer GH8HA01 and the pentamer Ac-GH8HA01Δ1Δ7. This latter peptide has a notably small mass (855.4 Da), nearing that of a small molecule and increasing its propensity for cellular permeability. These two peptides provide some of the most potent HDAC8 inhibition that has been determined while acting with a selectivety rarely surpassed by other, weaker HDAC isoform-selective inhibitors. While this is the second histone deacetylase and second ncAA warhead we have used with our PADLE technology to rapidly screen potent and selective inhibitors, we envision this approach can be applied to many other enzyme active sites with well defined interactions with pharmacophores or drug fragments. Much work has been done to expand the chemical toolbox of genetically encoded ncAAs.^43,44^ In combinations with these genetically encoded ncAAs, PADLE provides a means to conduct high throughput screening of phage-encoded peptides with minimal upfront costs or specialized techniques for the identification of potent and selective ligands for targets. We will continue to expand this tool kit for drug discovery.

## 4. MATERIALS AND METHODS

### 4.1 Expression and Purification of Apo-HDAC8 from *E. coli*

A pET14b plasmid coding for His-SUMO-HDAC8 was transformed into BL21(DE3pLysS) chemically competent cells. An overnight culture of the transformed cells was inoculated into autoinduction media (8 g/L of tryptone, 4 g/L yeast extract, 8 g/L NaCl, 4.56 g/L glycerol, 3.08 g/L KH_2_PO_4_, 3.10 g/L Na_2_HPO_4_•2H_2_O, 0.44 g/L MgSO_4_•7H_2_O, 0.45 g/L glucose, 1.2 g/L lactose) with chloramphenicol (34 µg/mL) and ampicillin (100 µg/mL) and let grown for 20 h at 24 °C. Cells were then harvested via centrifugation (8000 rcf, 10 min, 4 °C) and resuspended in 50 mL of lysis buffer (50 mM Tris pH 8.0, 150 mM KCl) with 50 µL of DNAse I (1 µg/mL final concentration) and with 5 mM dithiothreitol (DTT). The suspended cells were sonicated at 4 °C. The lysate was centrifuged to clarity (18000 rcf, 30 minutes 4 °C) and the supernatant was decanted to 5 mL of pre-equilibrated Ni-NTA resins. After incubation with agitation from 1 h, the resins were loaded to an empty chromatography cartridge and washed with 50 mL of wash buffer (lysis buffer plus 20 mM imidazole) and then His-SUMO-HDAC8 was eluted in elution buffer (lysis buffer plus 100 mM imidazole).

To the eluted His-SUMO-HDAC8 was added 5 mM DTT and 200 µg His SUMO Protease. This solution was dialyzed overnight at 4 °C in 35000 MWCO dialyzing tubing against wash buffer with 5 mM β-mercaptoethanol (BME) while cleavage of the His-SUMO tag occurred. The dialyzed solution was re-run over Ni-NTA pre-equilibrated with wash buffer, and then the flowthrough containing tag-free HDAC8 was collected. The resin was washed additionally with 5 mL more of wash buffer and the flowthrough was collected as well. A small portion of the solution containing tag-free HDAC8 was diluted to 10% glycerol, flash frozen in aliquots, and stored at -80 °C. The remaining portion was dialyzed against EDTA dialysis buffer (25 mM MOPS pH 7.5, 1 mM EDTA, 1 mM TCEP, 5 mM KCl) overnight and changed to fresh buffer once at 4 °C to remove bound metals in HDAC8 to make apo-HDAC8. The solution was then dialyzed against metal free buffer (25 mM MOPS pH 7.5, 1 mM TCEP, 5 mM KCl) overnight and changed to fresh buffer once at 4 °C. The apo-HDAC8 was flash frozen in aliquots and stored at -80 °C.

### 4.2 Expression and Purification of Apo-GST-HDAC8 from sf9 cells

First, a recombinant baculovirus was prepared with pAcGHLT-C coding for the expression of GST-HDAC8. Sf9 insect cells were seeded in 3 mL onto 60 mm tissue culture plates with 1.0 × 10^6^ Sf9 cells and incubated at 27 °C for 30 min, at which point they were determined to be 50-60% confluent. Cells were covered with 1 mL of fresh media. A transfection solution containing 2 *µ*g of plasmid, 2.5 µL of BaculoGold Bright Linearized baculovirus, and 5% PureFection in 100 µL was added to the cells and incubated at 27 °C for one week. Successful transfection was confirmed by the expression of GFP. 1 mL of the solution containing baculovirus was amplified in a T75 tissue culture flask in 9 mL of HyQ-SFX media with penicillin (100 µg/mL) and streptomycin (100 µg/mL) which was seeded with 2 × 10^6^ Sf9 cells, and baculovirus was expressed over 6 days at 27 °C. 10 mL of this solution was added to 2.5 × 10^6^ Sf9 cells in 500 mL of the same media and antibiotics and the cells were stirred at 120 rpm for 6 days at 27 °C. Supernatant was collected and stored at 4 °C for transduction.

Five flasks with 400 mL media with antibiotics were seeded with 1 × 10^6^ Sf9 cells each and 14 mL of GST-HDAC8 baculovirus solution was added to each. Flasks were shaken at 125 rpm at 27 °C for two days. Cells were harvested by centrifugation (4000 rpm for 5 minutes), and the supernatant was decanted. Cells were resuspended in 100 mL of lysis buffer (200 mM Tris pH 8.0, 0.15 M NaCl, 1% NP-40) and lysed via rocking at 4 °C for 2 h. Lysed cells were pelleted (16,000 rpm, 35 min, 4 °C), and the supernatant was added to 6 mL of fresh Ni-NTA resin, and incubated for 2 h at 4 °C. The resin was washed with 80 mL of wash buffer (20 mM Tris pH 7.5, 500 mM NaCl, 25 mM imidazole), and then the protein was eluted in 20 mL elution buffer (20 mM Tris pH 7.5, 150 mM NaCl, 250 mM imidazole). This elution was added to 0.75 mL of glutathione (GSH) agarose resin pre-equilibrated with elution buffer and incubated at 4 °C for 90 min. The resin was centrifuged (500 rcf, 5 min, 4 °C), the supernatant was decanted, and the resin was washed four times with 12 mL of modified phosphate buffered saline (high salt PBS, 500 mM NaCl, 2.7 mM KCl, 10 mM Na_2_HPO_4_, 1.8 mM KH_2_PO_4_, pH 7.4). GST-His-HDAC8 was eluted with GSH elution buffer (50 mM Tris pH 8.0, 15 mM GSH, 500 mL NaCl), flash frozen, and stored in single-use aliquots at -80 °C.

### 4.3 Expression and Purification of Aoda-Containing Phages

Electrocompetent TOP10 *E. coli* cells containing pEVOL-CloDF-PylT-AcKRS (pEVOL-PylT-AcKRS with a CloDF origin of replication) and M13KO7(pIII^-^) (M13KO7 helper phage with a nonsense TAA mutation in gIII) were transformed with pADL(NNK)_7_pIII in which a TAG-enriched (NNK)_7_ library was appended to the 5’ end of the gIII gene. Transformed cells were grown in 2xYT with ampicillin (100 µg/mL), chloramphenicol (34 µg/mL), and kanamycin (25 µg/mL) and stored in one-tenth volume 20% glycerol aliquots at -80 °C. These aliquots were diluted into 1.1 L of fresh 2×YT media with ampicillin, chloramphenicol, and kanamycin, and inoculated with stored cells to OD_600_ = 0.1. Cells were incubated at 37 °C until OD_600_ = 0.5-0.8, at which point 1 mM IPTG and 0.2% arabinose were added to induce phage expression. 100 mL of cells were added to a sterile flask to express the negative control, while 10 mM of racemic Aoda hydrochloride was added to the remaining 1 L of cells. Phages were expressed at 30 °C for 16 h. Cells were pelleted via centrifugation (4,000 rcf, 20 min, 4 °C), and the supernatant was poured into 5× precipitation buffer (2.5M NaCl, 20% PEG-8000). The phages were precipitated at 4 °C for two h. Phages were pelleted via centrifugation (10,000 rcf, 25 min, 4 °C), and the supernatant was discarded. Phages were then resuspended in 30 mL of phage binding buffer (50 mM HEPES pH 8.0, 137 mM NaCl, 2.7 mM KCl, 1 mM MgCl_2_), centrifuged to clarify (4,000 rcf, 20 min, 4 °C), 7.5 mL of 5× precipitation buffer was added, and phages were precipitated at 4 °C. Phages were once again pelleted via centrifugation (10,000 rcf, 25 min, 4 °C), the supernatant decanted, and phages were dissolved in 1 mL of phage binding buffer. Phages were then centrifuged to clarity (14,000 rcf, 10 min, room temperature), and the phage solution was heat shocked at 65 °C for 15 minutes to kill any remaining cells. Final phages were stored at 4 °C until used for selection.

To titer the purified phage solution, ER2738 *E. coli* cells were grown in 2×YT with tetracycline (10 µg/mL) at 37 °C until OD_600_ = 0.5-0.8. 10 µL of phages were diluted into 90 µL of 2×YT, and serially diluted ten-fold. 10 µL of diluted phages were added to 90 µL of ER2738 and incubated at 37 °C for 50 min. 10 µL of the infected cultures were spotted onto LB agar with ampicillin (100 µg/mL) and tetracycline (10 µg/mL) and incubated at 37 °C overnight. The colonies in each spot were counted and this number was used to determine the number of colony forming units (cfu) in the original phage solution.

### 4.4 Affinity Selection of Aoda-Containing Phages Against GST-HDAC8

50 µL of Pierce GSH magnetic agarose beads (Thermo Fisher Scientific) were washed three times with 1 mL of phage binding buffer (50 mM HEPES pH 8.0, 137 mM NaCl, 2.7 mM KCl, 1 mM MgCl_2_). 1 mL of binding buffer containing 18 µg of GST-HDAC8 was added to the beads, and the beads were incubated at room temperature for 30 min under slow rocking. The supernatant was removed, and beads were washed three times with 1 mL of phage binding buffer. The beads were then incubated with 1x blocking buffer (binding buffer with 1% BSA and 0.1% Tween-20) for 30 min under slow rocking. To 1 mL of phages were added 250 µL of 5× blocking buffer (binding buffer with 5% BSA and 0.5% Tween-20) and phages were blocked for the same amount of time. After 30 min, the blocking buffer was removed from the magnetic beads and the blocked phages were added, and the phages were incubated with the beads for 30 min under slow rocking. Phages were then removed, and the beads were washed with 1 mL of wash buffer (blocking buffer with 0.1% Tween-20). For rounds 1 and 2, beads were washed three total times; for round 3, four washes were performed. Beads were transferred to a fresh tube after the first wash. 100 µL of elution buffer (50 mM glycine pH 2.2) were added to the beads, which were gently agitated for 15 min. The elution solution was removed from the beads and neutralized into 50 µL of neutralization buffer (1 M Tris pH 8.0). 25 mL of 2×YT with tetracycline (10 µg/mL) were inoculated with ER2738 cells and grown to OD_600_ = 0.5-0.8. 5 mL of ER2738 cells were removed, and 140 µL of the eluted phage solution was added to the remaining 20 mL. Cells were transfected for 50 min at 37 °C, and then centrifuged (4,000 rcf, 10 min, 4 °C). The supernatant was discarded, and the cells were resuspended in 500 mL of 2xYT with tetracycline (10 µg/mL) and ampicillin (100 µg/mL) and grown overnight at 37 °C. Amplified transfected cells were stored in one-tenth volume aliquots in 20% glycerol at -80 °C and used to express phages for the next proceeding round. The remaining 10 µL of eluted phages were used to quantify the total elution with the remaining ER2738 cell culture.

For the 2nd and 3rd round of affinity selection, a negative control was added to the procedure. An additional 50 µL of Pierce GSH magnetic agarose beads was washed three times with 1 mL of binding buffer, and to these beads was added 1 mL of binding buffer with 8 µg of GST, and the beads for negative selection were incubated for 30 min. These beads were washed three times with 1 mL of binding buffer, and then blocked with 1 mL of 1x blocking buffer for 30 min. After both sets of beads and the phages were blocked, the blocking solution was removed from the negative selection beads, and phages were added to these beads, and incubated under slow rocking for 30 minutes. The blocking solution was then removed from the magnetic beads containing GST-HDAC8, and the supernatant from the negative selection beads were removed and directly added to the positive selection beads, incubating for 30 min under slow rocking. The affinity selection followed the same procedure as above for all proceeding steps.

### 4.5 Synthesis and Affinity Characterization

The synthesis of substrates and peptides and affinity characterization of synthesized peptides are provided in the Supplementary Information.

## 5. AUTHOR CONTRIBUTIONS

**Jared S. Morse:** Investigation (leading), writing (leading), review (leading) and editing (leading). **Yan Sheng:** Investigation (supporting). **J. Trae Hampton:** Investigation (supporting) and writing (supporting). **Sukant Das:** Investigation (supporting). **Yugendar R. Alugubelli:** Investigation (supporting). **Peng-Hsun Chase Chen:** Investigation (supporting). **Kai S. Yang:** Investigation (supporting). **Shiqing Xu:** Conceptualization (supporting). **Carol A. Fierke:** Conceptualization (supporting). **Wenshe Ray Liu:** Conceptualization (leading), funding acquisition (leading), project administration (leading), and writing (leading).

## ACKNOWLEDGEMENTS

This work was support in part by National Institutes of Health (Grants R35GM145351 to W.R.L.), Cancer Prevention and Research Institute of Texas (a postdoctoral fellowship from Grant RP210043 to support K.S.Y.), Welch Foundation (Grant A-1715 to W.R.L.), the Texas A& M X-Grants Mechanism, and the Texas A& M Chancellor EDGES Fellowship. We would like to acknowledge Dr. Pingwei Li at Texas A& M University for assistance in expressing GST-HDAC8 in insect cells and Dr. Franz-Josef Meyer-Almes at University of Applied Sciences Darmstadt for providing the pET14b *E. coli* expression vector for His-SUMO-HDAC8.

## CONFLICT OF INTEREST

The authors declare no conflicts of interest.

## Notes

### Competing Interest Statement

The authors have declared no competing interest.

