## Supplementary Information for "Phage-Assisted, Active Site-Directed Ligand Evolution of a Potent and Selective Histone Deacetylase 8 Inhibitor"

For

### Supplementary Methods

#### 1. Protein Expression

Expression of Glutathione-S-Transferase

BL21(DE3) chemically competent cells were transformed with the pGEX-4T-3 plasmid. These cells were grown in 1L 2YT with ampicillin (100 µg/mL) at 37 °C until $OD_{600}$ = 0.5-0.8, at which point glutathione-S-transferase (GST) expression was induced with 0.1 mM isopropyl β-D-1-thiogalactopyranoside (IPTG) for 5 hours. Cells were centrifuged (4000 rpm, 10 min) and resuspended in 50 mL of phosphate buffered saline (PBS, 137 mM NaCl, 2.7 mM KCl, 10 mM $Na_2HPO_4$, 1.8 mM $KH_2PO_4$, pH 7.4) with 0.1% Tween-20 and the cOmplete protease inhibitor cocktail tablets (Roche). Cells were lysed via sonication, cellular debris was pelleted (16000 rcf, 35 min, 4 °C), the lysate was added to 1.5 mL of pre-equilibrated GSH agarose resin and incubated at 4 °C for 90 min. The resin was centrifuged (500 rcf, 5 min, 4 °C), washed four times with 25 mL of cold PBS, and then GST was eluted with GSH elution buffer, dialyzed into storage buffer (50 mM Tris, 150 mM NaCl, pH 8.0), flash frozen, and stored in aliquots at -80 °C.

Expression of sfGFP-N134TAG with KetoK Hydrochloride

Plasmids pEVOL-PylT-AcKRS and pBAD-sfGFP-N134(TAG), which contains the gene encoding superfolder green fluorescent protein (sfGFP) with an amber mutation at position N134, were co-transformed into chemically competent TOP10 *E. coli* cells. These cells were inoculated into 6 mL of 2YT with ampicillin (100 µg/mL) and chloramphenicol (34 µg/mL) and grown to $OD_{600}$ = 0.5-0.6 at 37 °C. 0.2% arabinose and 1 mM IPTG were added, and the media was split into two sterile tubes. To one, 4 mM of racemic KetoK hydrochloride was added, and sfGFP was expressed at 37 °C for 16 h. Cells were then pelleted, resuspended in lysis buffer (50 mM Tris pH 8.0, 150 mM NaCl, 0.1 mM PMSF) and lysed via three consecutive freeze-thaw cycles. Lysate was clarified via centrifugation (16000 rpm, 30 min, room 4 °C), and the supernatant was added to pre-equilibrated Ni-NTA resin (50 µL), incubated for 30 min, and then the supernatant was discarded, the resin was washed with 5 mL of wash buffer (50 mM Tris pH 8.0, 150 mM NaCl, 20 mM imidazole), and then sfGFP was eluted in elution buffer (wash buffer with 250 mM imidazole). A small aliquot of sfGFP expressed with and without KetoK was analyzed via 12% SDS-PAGE. The remaining solution of the KetoK-containing sfGFP was desalted into high purity water using 0.5 mL Zeba spin desalting columns 7K MWCO (Thermo Fisher Scientific). The desalted solution was analyzed via electrospray ionization (ESI) mass spectrometry.

#### 2. Preparing and Analyzing the Enriched Library for Next Generation Sequencing

Amplicons were generated by PCR using the following two primers purchased from IDT DNA Inc.: 1) 5'- TCGTCGGCAGCGTCAGATGTGTATAAGAGACAGGCCCAGCCGGC-CATG -3' and (2) 5'- GTCTCGTGGGCTCGGAGATGTGTATAAGAGACAGCGGCCGCTTT-CGCCGC -3'. The following PCR conditions were used:

**Table S1:** PCR Conditions

| Solution | Volume (µL) | Final Concentration |
|---|---|---|
| 5x Phusion GC Buffer | 10 | 1x |
| 10 mM dNTP | 1 | 0.2 mM |
| Sample DNA | 1 | 0.71 ng/µL |
| Forward Primer | 0.5 | 1 µM |
| Reverse Primer | 0.5 | 1 µM |
| High Purity Water | 36 | |
| Phusion HiFidelty Polymerase | 1 | 2U |

**Table S2:** PCR Cycles

<table>
<tr><th>Temperature (°C)</th><th>Time (s)</th><th>Cycles</th></tr>
<tr><td>98</td><td>60</td><td></td></tr>
<tr><td>98</td><td>10</td><td rowspan="4">30</td></tr>
<tr><td>55</td><td>10</td></tr>
<tr><td>69</td><td>30</td></tr>
<tr><td>72</td><td>30</td></tr>
<tr><td>12</td><td>Hold</td><td></td></tr>
</table>

The amplicons were purified and extracted from a 3% agarose gel according to the GenCatch gel extraction kit. These amplicons were sent to the Genomics and Bioinformatics center at Texas A&M University, where the enriched libraries were then sequenced using Illumina MiSeq paired end sequencing (2 x 150 bp). Sequences were analyzed in R. Reads containing the two primer regions flanking the library sequence were then spliced out of the initial sequences of both forward and reverse reads. Paired-end processing[1] was adapted to the R platform—any reads containing more than one mismatch in the primer regions or any mismatch in the library region were discarded. A matrix was created to analyze the amino acid coded for in each position of the randomized region of the library, treating the amber codon as a 21st amino acid. Relative enrichment was measured as the relative rate of appearance of each unique amino acid sequence occurred in round 3 minus its relative rate of appearance in round 1, divided by the relative rate of appearance of the same sequence in round 1. An enrichment score > 0 corresponds to a positive enrichment throughout the selection experiment, an enrichment score < 0 corresponds to a loss of the prevalence of the sequence throughout the selection experiment, and an enrichment score of 0 corresponds to no change in prevalence of the individual sequence in the enriched library through iterative rounds of selection.

#### 3. Synthesis

The Synthesis of Aoda

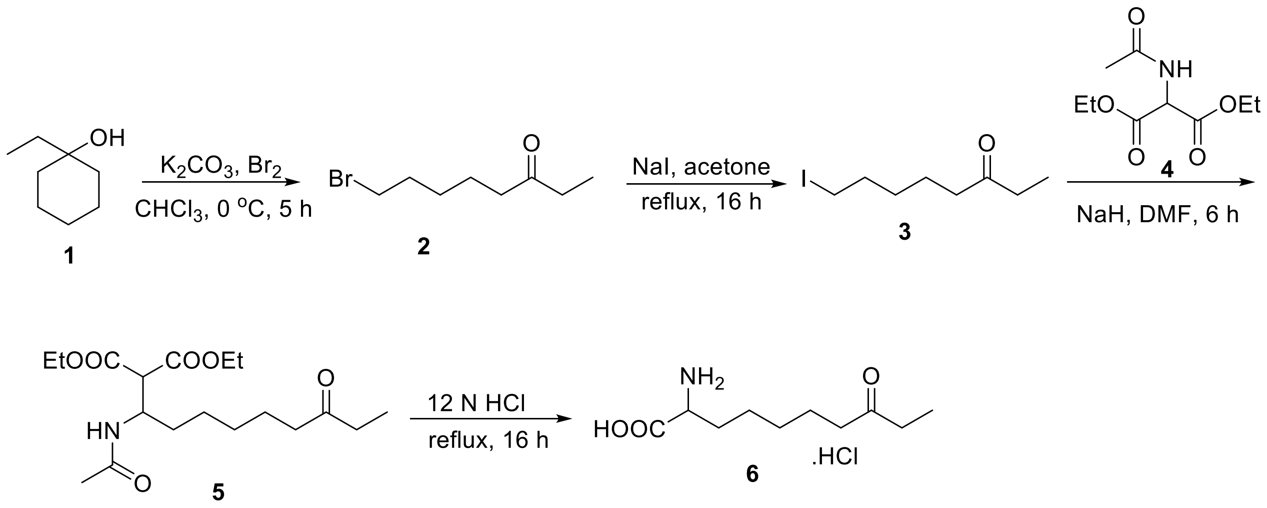


**Scheme S1**: A revised synthetic route for Aoda (**6**)

8-bromooctan-3-one (**2**): To a solution of cyclic tertiary alcohol **1** (1 g, 7.8 mmol) in $CHCl_3$ (20 mL) was added $K_2CO_3$ (6.4 g, 46.8 mmol) at 0 °C. The mixture was stirred for 10 min, and bromine (2.0 mL, 39 mmol) was then added dropwise. The reaction mixture was stirred at 0 °C for 5 h. The mixture was quenched with sat. $Na_2S_2O_3$ solution (20 mL) and extracted with EtOAc (2×50 mL).

The combined organic layers were washed with brine, dried over $Na_2SO_4$ and concentrated, which was used in the next step without purification.

8-iodooctan-3-one (**3**): A mixture of **2** (1 g, 4.85 mmol), NaI (2.1 g, 14.5 mol) in acetone (20 ml) was heated at reflux for 18 h. The resulting mixture was concentrated under reduced pressure and was then taken up in $Et_2O$ (20 ml), filtered, and the filter cake was washed with $Et_2O$ (10 mL). The combined $Et_2O$ filtrates were washed with $H_2O$ (2x5 mL), sodium thiosulfate solution (5 mL) and brine (2 mL). The resulting solution was dried over $Na_2SO_4$ and concentrated under reduced pressure to yield the desired material (1 g) which was used in the next step without purification.

Diethyl 2-(1-acetamido-7-oxononyl)malonate (**5**): To a stirred solution of diethyl 2-acetamidomalonate **4** (0.6 g, 2.7 mmol) in anhydrous DMF (10 mL) was added NaH (71 mg, 2.9 mmol) at 0 °C. The mixture was stirred for 10 min, and iodo ketone **3** (736 mg, 2.9 mmol) was then added dropwise. The reaction mixture was stirred at rt for 6 h. After this $Et_2O$ was added to the mixture and the solution was washed with $H_2O$ (3x15 ml), 0.1 M HCl (15 ml), sat. $NaHCO_3$ (15 ml) and brine (15 ml). The organic layer was dried over $Na_2SO_4$ and concentrated and purified by silica gel chromatography (20-50 % EA/Hexane) to provide the compound (800 mg, 81%).

2-Amino-8-oxodecanoic acid (**6**): A mixture of **5** (0.7 g, 1.96 mmol), and 12 N HCl (10 ml) was heated at reflux for 18 h. The reaction mixture was concentrated and triturated with ether to provide the desired product as white solid (450 mg, 84%). $^1$H NMR (400 MHz, DMSO-$d_6$) δ 8.49 (s, 3H), 3.80 (t, $J$ = 6.0 Hz, 1H), 2.40 (q, $J$ = 7.4 Hz, 4H), 1.85 – 1.68 (m, 2H), 1.42 (dp, $J$ = 11.7, 6.9 Hz, 3H), 1.33 – 1.13 (m, 3H), 0.90 (t, $J$ = 7.3 Hz, 3H).

The synthesis of Fmoc-Aoda

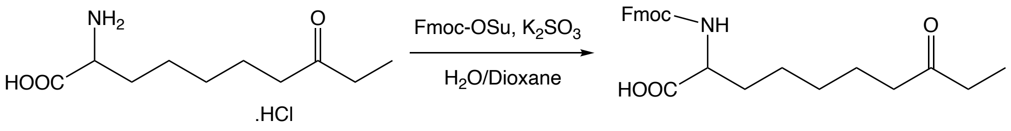


**6** **7**

**Scheme S2: A synthetic route for Fmoc-Aoda**

A suspension of potassium carbonate (2.19 g, 15.8 mmol) and **6** (1.0 g, 4.2 mmol) in DI water (45 ml) was stirred at 0 °C. A solution of 9-fluorenylmethyl N-succimidyl carbonate (1.42 g, 4.2 mmol) in dioxane (40 mL) was added dropwise to the stirred suspension over the course of 1 hour. The reaction was allowed to warm to room temperature and stirred overnight. The solution was concentrated under reduced pressure to remove dioxane, and then acidified with 20% citric acid (aq.) until pH < 4. The reaction was extracted with ethyl acetate (EtOAc, 4 x 25 mL). The organic layer was dried over $Na_2SO_4$, filtered, and concentrated under reduced pressure. Crude product was purified via flash chromatography (methanol/dichloromethane, 0%-6%) to give Fmoc-KetoK-OH (1.2 g, 67%) as a white solid. 1H NMR (400 MHz, DMSO-$d_6$) δ 7.89 (d, 2H), 7.72 (d, 2H), 7.62 (d, 1H), 7.42 (t, 2H), 7.32 (t, 2H), 4.29-4.20 (m, 3H), 3.94-3.88 (m, 1H), 2.42-2.37 (m, 4H), 1.71-1.16 (m, 8H), 0.90 (t, 3H).

<u>The Synthesis of *L*-*N*-Fmoc-2-amino-8-[*O*-(4-methoxyphenylmethyl)hydroxyamino]-8-oxooctanoic acid (Fmoc-Asuha(NHOPMB-OH)</u>

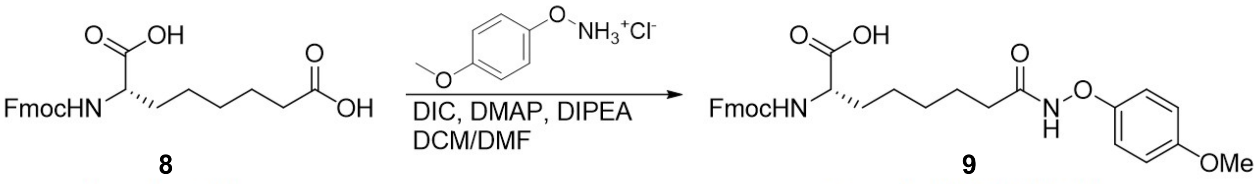


**8** **9**

Fmoc-Asu-OH Fmoc-Asu(NHOPMB)-OH

**Scheme S3: A synthetic route for Fmoc-Asu(NHOPMB)-OH**

To a solution of **8** (0.40 g, 0.97 mmol) in DCM (20 mL) and DMF (1 mL) was added 1-ethyl-3-(3-dimethylaminopropyl)carbodiimide (0.143 g, 0.92 mmol), and the solution was stirred on ice. DMAP (11 mg, 92 μmol) was added, and then a solution of O-(4-methoxybenzyl)hydroxylamine hydrochloride (0.176 g, 0.93 mmol) and DIPEA (322 μL, 1.85 mmol) in DCM (6 mL) was added drop-wise to the stirred solution. The reaction was allowed to warm to room temperature and stirred overnight. The solution was washed with 1 N HCl (3 x 15 mL), brine (15 mL), dried over $MgSO_4$, filtered, and concentrated under reduced pressure to remove solvent. After flash chromatography (EtOAc/hexanes [both with 5% acetic acid), 30% to 70%), **9** (190 mg, 38%) was obtained as a white solid. 1H NMR (400 MHz, DMSO-$d_6$) δ 12.14 (bs, 1H), 10.85 (s, 1H), 7.89 (d, 2H), 7.72 (d, 2H), 7.61 (d, 1H), 7.42 (t, 2H), 7.34-7.28 (m, 4H), 6.92 (d, 2H), 4.69 (s, 2H), 4.29-4.20 (m, 3H), 3.94-3.88 (m, 1H), 3.74 (s, 3H), 1.95-1.90 (m, 2H), 1.72-1.43 (m, 4H), 1.36-1.16 (m, 4H).

<u>The Synthesis of HDAC8 Substrate</u>

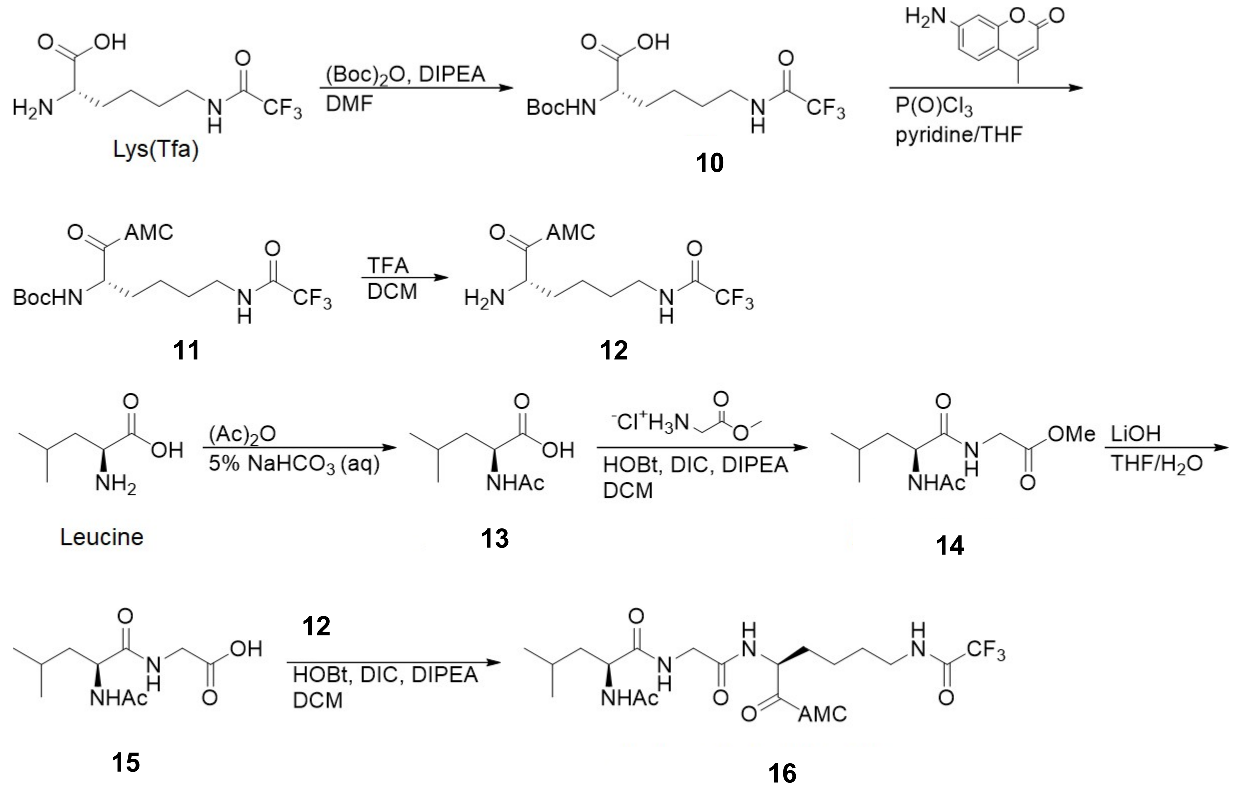


**Scheme S4: A synthetic route for HDAC8 substrate**

The synthetic route shown in Scheme S4 was adapted from several literature procedures.[2-4]

$N^{\varepsilon}$-trifluoroacetyllysine (Lys(Tfa), 1.0 g, 4.1 mmol) was suspended in dry DMF (11 mL) under an atmosphere of nitrogen, and then Boc-anhydride (1.04 mL, 4.53 mmol) and DIPEA (1.4 mL, 8.2 mmol) were added, and the reaction was stirred overnight. The reaction was diluted with 1 N HCl (25 mL), extracted with EtOAc (2 x 20 mL), and the combined organic layers were washed with 1 N HCl (25 mL), brine (25 mL), dried over $MgSO_4$, filtered, and concentrated under reduced pressure to give **10** (285 mg, 20%) as a solid. Product was used directly in the next step.

**10** (285 mg, 0.833 mmol) and 7-amino-4-methylcoumarin (AMC, 171 mg, 0.976 mmol) was dissolved in 16 mL of dry THF and cooled on ice. To this solution was added pyridine (0.72 mL, 8.9 mmol), and the reaction was stirred for 30 min. Then, phosphoryl chloride (0.29 mL, 3.11 mmol) was added dropwise, and the reaction was stirred on ice for 2.5 h followed by 2 h at room temperature. Then, the reaction was poured into a 1:1 mixture of ice and saturated sodium bicarbonate (20 mL), concentrated under reduced vacuum to approximately 20 mL, and then extracted with EtOAc (3 x 25 mL). The combined organic layers were washed with 1 N HCl (2 x 30 mL), saturated sodium bicarbonate (30 mL), brine (30 mL), dried over $MgSO_4$, filtered, and then concentrated under reduced pressure to give **11** (310 mg, 75%) as an orange solid and used directly in the next step.

**11** (98 mg, 0.20 mmol) was dissolved in DCM (2 mL) and trifluoroacetic acid (TFA, 1 mL), and stirred for 4 h at room temperature. The reaction was concentrated under reduced pressure to give **12** (100 mg, quant.). 1H NMR (400 MHz, DMSO-$d_6$) δ 10.85 (s, 1H), 9.39 (t, 1H), 8.25 (d, 3H), 7.81-7.76 (m, 2H), 7.46 (dd, 1H), 6.38 (d, 1H), 3.94 (m, 1H), 3.22-3.14 (m, 2H), 2.42 (d, 3H), 1.88-1.80 (m, 2H), 1.56-1.47 (m, 2H), 1.40-1.31 (m, 2H).

Leucine (2.0 g, 15.2 mmol) was suspended in a 5% solution of sodium bicarbonate (43 mL) and cooled on ice. Acetic anhydride (1.74 mL, 18.4 mmol) was added dropwise over 45 min, and the reaction was allowed to warm to room temperature as it stirred for 4 h. Reaction was acidified to pH 2 with 6 N HCl, and product was precipitated on ice. The precipitate was filtered, washed with cold water (2 x 10 mL), and dried under vacuum overnight, giving Ac-Leu-OH (**13**, 1.37 g, 52%), and was used directly in the next step.

**13** (526 mg, 3.00 mmol) and hydroxybenzotriazole (HOBt, 436 mg, 3.22 mmol) were suspended in dry DCM (5 mL) and cooled on ice. Diisopropylcarbodiimide (DIC, 0.51 mL, 3.7 mmol) was added and stirred for 5 min. Then, a suspension of glycine hydrochloride methyl ester (426 mg, 3.39 mmol) and DIPEA (0.56 mL, 3.2 mmol) in DCM (7 mL) was added to the reaction, and the reaction was allowed to warm to room temperature as it stirred overnight. Then, 3 mL of methanol was added to solubilize any solids in the reaction flask. 1 N HCl (30 mL) was added, and the organic layer was removed. EtOAc (2 x 25 mL) was used to back extract the aqueous layer. The combined organic layers were washed with 1 N HCl (30 mL), and this aqueous wash was back extracted with EtOAc (2 x 25 mL). The combined organic layers were washed with saturated sodium bicarbonate (30 mL), and this aqueous layer was back extracted with EtOAc (2 x 25 mL). The combined organic layers were washed with brine (30 mL), dried over $MgSO_4$, filtered, and concentrated under reduced pressure. The resulting crude Ac-Leu-Gly-OMe (**14**, 200 mg, 27%) was obtained as white solid and used directly in the next step.

**14** (200 mg, 0.82 mmol) was dissolved in dry THF (4 mL). To this was added 1 M LiOH (4 mL), and the reaction was stirred at room temperature for 1 h. The reaction was concentrated under reduced pressure to remove THF, and then 2 N HCl was added until pH = 2. The aqueous layer was extracted with EtOAc (8 x 5 mL). The combined organic layers were dried over $MgSO_4$, filtered, and rotovaped, giving Ac-Leu-Gly-OH (**15**, 141 mg, 75%) as a solid. The product was used without further purification in directly in the next step.

**15** (23 mg, 0.10 mmol) and HOBt (14.2 mg, 0.105 mmol) were suspended in dry DCM (1.3 mL) under nitrogen. DIC (18 μL, 0.11 mmol) was added, and the reaction was stirred for 25 min. Amine **S7** (50 mg, 0.097 mmol) and DIPEA (34 μL, 0.20 mmol) were dissolved in dry DCM (0.8 mL) and added slowly to the reaction. The reaction was stirred at room temperature for 2 h, and then brine (5 mL) and DCM (4 mL) was added, and the layers separated. The aqueous layer was back extracted with DCM (3 x 4 mL). The combined organic layers were washed with 1 N HCl (2 x 5 mL), saturated sodium bicarbonate (5 mL), and then the sodium bicarbonate solution was back extracted with DCM (2 x 4 mL). The combined organic layers were dried over $MgSO_4$, filtered, and concentrated under reduced pressure. The crude product was purified via flash chromatography (MeOH/DCM, 4% to 10% MeOH), giving the resulting HDAC8 substrate Ac-

LGK(Tfa)-AMC (**16**, 28 mg, 47%) as an off-white solid. 1H NMR (400 MHz, DMSO-$d_6$) δ 10.38 (s, 1H), 9.41 (t, 1H), 8.32 (t, 1H), 8.07 (dd, 2H), 7.80 (d, 1H), 7.74 (d, 1H), 7.53 (dd, 1H), 6.28 (d, 1H), 4.39 (m, 1H), 4.22 (m, 1H), 3.73 (m, 2H), 3.17 (m, 2H), 2.41 (d, 3H), 1.85 (s, 3H), 1.80-1.20 (m, 9H), 0.87 (m, 6H).

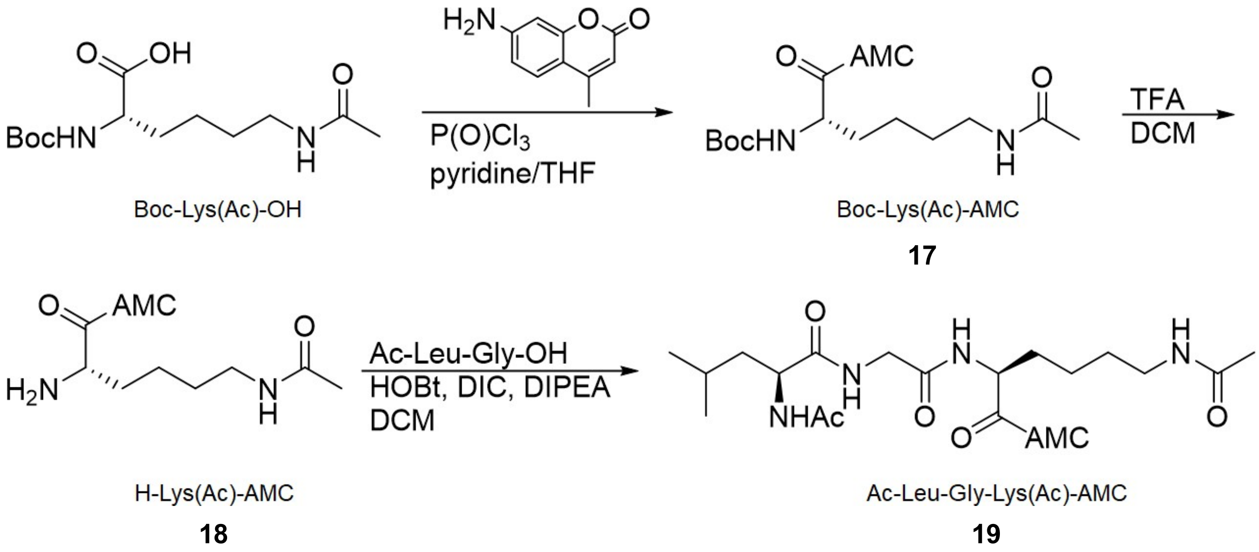


**Scheme S5:** A synthetic route for HDAC1, 2, and 6 substrate

The synthetic route shown in Scheme S4 was adapted from several literature procedures.[2,3,5]

Boc-Lys(Ac)-OH (0.576 g, 2.00 mmol) was dissolved in dry DMF (10 mL) under nitrogen, and cooled on ice. HATU (0.989 g, 2.60 mmol) and DIPEA (1.4 mL, 8.0 mmol), and the reaction was stirred for 10 min. Then, AMC (0.350 g, 2.00 mmol) was added, and the reaction was warmed to 60 °C and stirred overnight. Saturated sodium bicarbonate (5 mL) was added, and then water was added to resolubilize the precipitate (50 mL). EtOAc (3 x 15 mL) was used to extract, and then the combined organic layers were washed with 1 N HCl (2 x 30 mL), brine (30 mL), dried over $MgSO_4$, filtered, and concentrated under reduced pressure. After flash chromatography (MeOH/DCM, 3% to 7% MeOH), the product Boc-Lys(Ac)-AMC (**17**, 250 mg, 28%) was obtained. 1H NMR (400 MHz, $CDCl_3$) δ 9.24 (s, 1H), 7.66 (d, 1H), 7.51 (m, 2H), 6.18 (d, 1H), 5.83 (m, 1H), 5.32 (d, 1H), 4.25 (m, 1H), 3.29 (m, 2H), 2.40 (d, 3H), 2.01 (s, 3H), 1.76-1.38 (m, 15H)

**17** (100 mg, 0.224 mmol) was deprotected in the same manner as Boc-Lys(Tfa)-AMC. The resulting H-Lys(Ac)-AMC (**18**, 103 mg, quant.) was used directly without purification.

**18** (100 mg, 0.22 mmol) was coupled to Ac-Leu-Gly-OH using the same procedure to synthesize Ac-Leu-Gly-Lys(Tfa)-AMC. After flash chromatography (MeOH/DCM, 4%-15%), the substrate for HDAC 1, 2, and 6, Ac-LGKAc-AMC (**19**, 23 mg, 19%) was obtained. 1H NMR (400 MHz, $CDCl_3$) δ 10.37 (s, 1H), 8.32 (t, 1H), 8.09 (d, 1H), 8.00 (d, 1H), 7.82-7.77 (m, 2H), 7.73 (d, 1H), 7.53 (dd, 1H), 6.27 (m, 1H), 4.37 (m, 1H), 4.22 (q, 1H), 3.73 (m, 2H), 3.00 (q, 2H), 2.40 (d, 3H), 1.84 (s, 3H), 1.77 (s, 3H), 1.67-1.55 (m, 3H), 1.48-1.22 (m, 6), 0.86 (dd, 6H).

Solid Phase Peptide Synthesis of Selected Peptides

Peptides were synthesized on ProTide Rink amide low loading resin (CEM #R002) using an automated Liberty Blue Peptide Synthesizer with an HT12 attachment. Fmoc-amino acids were deprotected using 20% piperidine in DMF with a two-step microwave cycle: 75 °C for 15 seconds, then 90 °C for 50 seconds. Free Fmoc-amino acids (5 eq.) were then coupled with 0.25 M Oxyma and 0.125 M DIC in DMF using the Liberty Blue's standard microwave coupling cycle: 75 °C for 15 seconds, then 90 °C for 110 seconds. Aoda was installed using racemic Fmoc-Aoda-OH (2.5 equivalents), and Asuha was installed using Fmoc-Asu(NHOPMB)-OH (5 eq.). the *N*-terminal acetylation was done with 3 mL of 25% acetic anhydride in DCM for 5 min at room temperature. Peptides were cleaved from resin by agitating for 3 h in 2 mL of cleavage solution (95% TFA, 5% DCM). The products were filtered and concentrated by bubbling nitrogen and then dried by removing solvent under vacuum overnight. The peptides were dissolved in 450 µL of DMF and purified by reverse-phase semi-preparative HPLC (20%-60% acetonitrile in water with 1% formic acid, 40 minutes, 18 mL/min, PDA) using a Shimadzu Shim-pack GIS C18 column 10 µm (25 mm x 250 mm, Shimadzu #227-30115-04).

#### 4. Inhibitor Validation

#### $IC_{50}$ Determination with Reconstituted HDAC8

Apo-HDAC8 was reconstituted with $Zn(NO_3)_2$ (1 equivalent) in HEPES assay buffer (25 mM HEPES pH 8.0, 3 mM KCl, 137 mM NaCl, 0.1 mg/mL BSA, 0.0002% Tween-20) for 1 h at 4 °C. Inhibitors were dissolved in DMSO and serially diluted in DMSO/water (2.5% DMSO), and then preincubated at room temperature with reconstituted HDAC8 (0.5 nM in assay buffer) for 10 min. 50 μM HDAC8 substrate (Ac-LGK(Tfa)-AMC) was then added, and the reaction was incubated for 60 min at 30 °C. A freshly prepared quench solution (1 volume equivalent, 2 μM TSA, 2 mg/mL trypsin in assay buffer) was added to stop the reaction and develop the fluorescent signal. The plate was shaken for 10 min, and then the fluorescence at 360 nm/460 nm was measured.

#### $IC_{50}$ Determination with HDAC1, 2, and 6

HDACs 1, 2, and 6 (BPS Bioscience) were diluted in HEPES assay buffer (0.8 nM HDAC1, 2.4 nM HDAC2, and 0.8 nM HDAC6). Inhibitors were dissolved in DMSO and serially diluted in DMSO/water (2.5% DMSO), and then preincubated at room temperature with the HDAC for 10 min. 20 μM HDAC1, 2, and 6 substrate (Ac-LGKac-AMC) was then added, and the reaction was incubated for 30 min at 37 °C. A freshly prepared quench solution was added (1 volume equivalent) to stop the reaction and develop the fluorescent signal. The plate was shaken for 10 min, and then the fluorescence at 360 nm/460 nm was measured.

#### Supplementary Figures

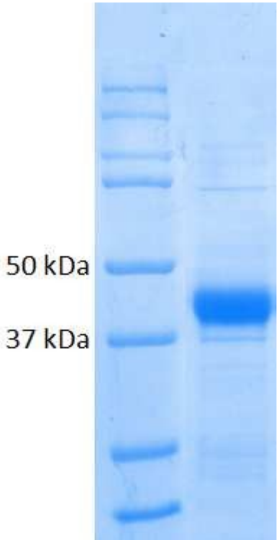


**Figure S1:** Expression of HDAC8 in *E. coli*. Expected molecular weight: 42 kDa.

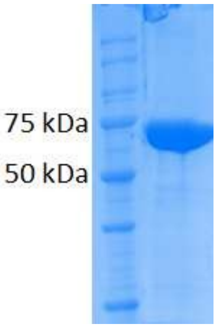


**Figure S2:** Expression of GST-His-HDAC in sf9 cells. Expected molecular weight: 70 kDa.

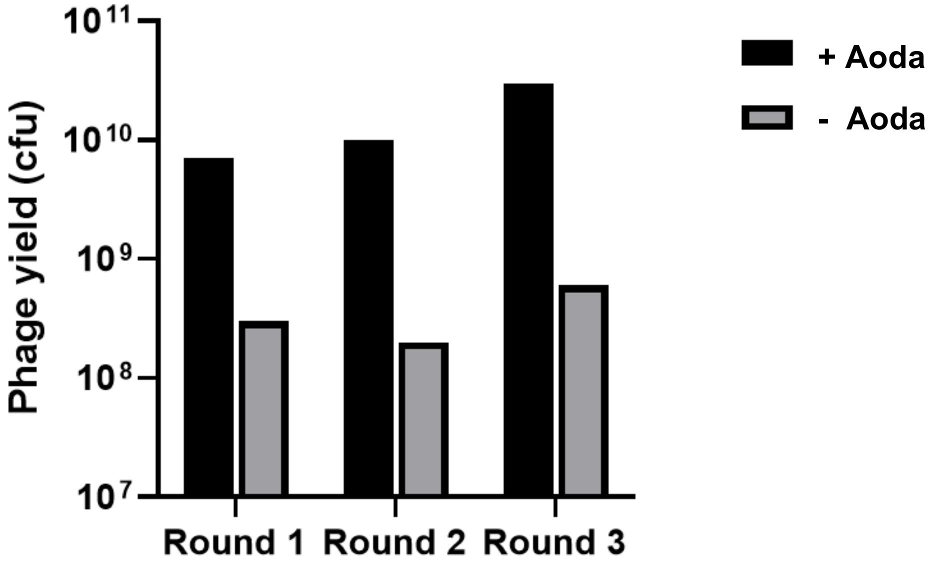


**Figure S3:** Phage yields in the presence and absence of 10 mM Aoda·HCl through three rounds of selection. Through three rounds of selection against GST-His-HDAC8, sequences containing amber codons are enriched relative to phages coding for canonical sequences.

#### HPLC/MS Of Synthesized Peptides

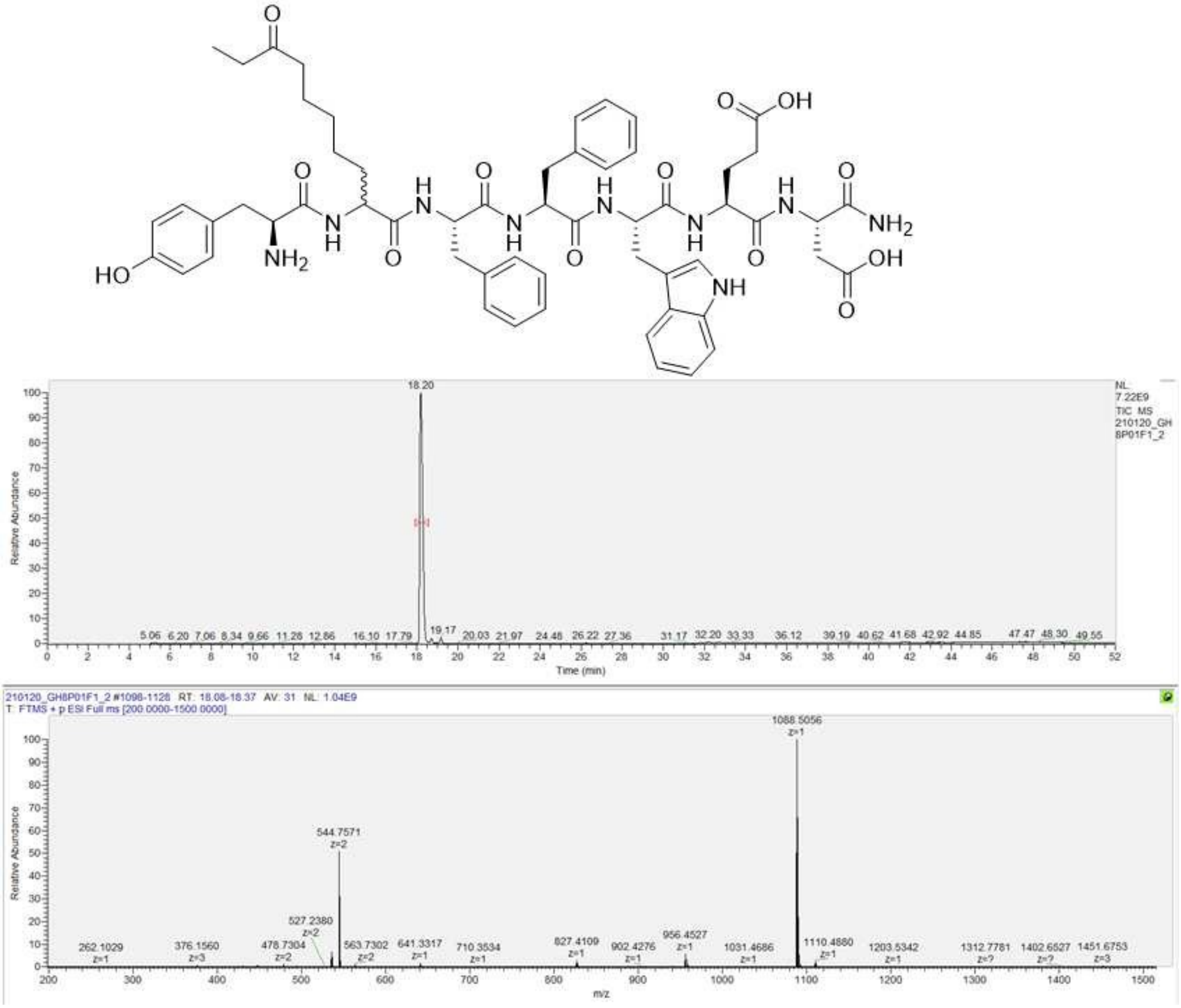


**Figure S4:** HPLC/MS of GH8P01F1. Calculated M+H$^+$: 1088.5088 Da.

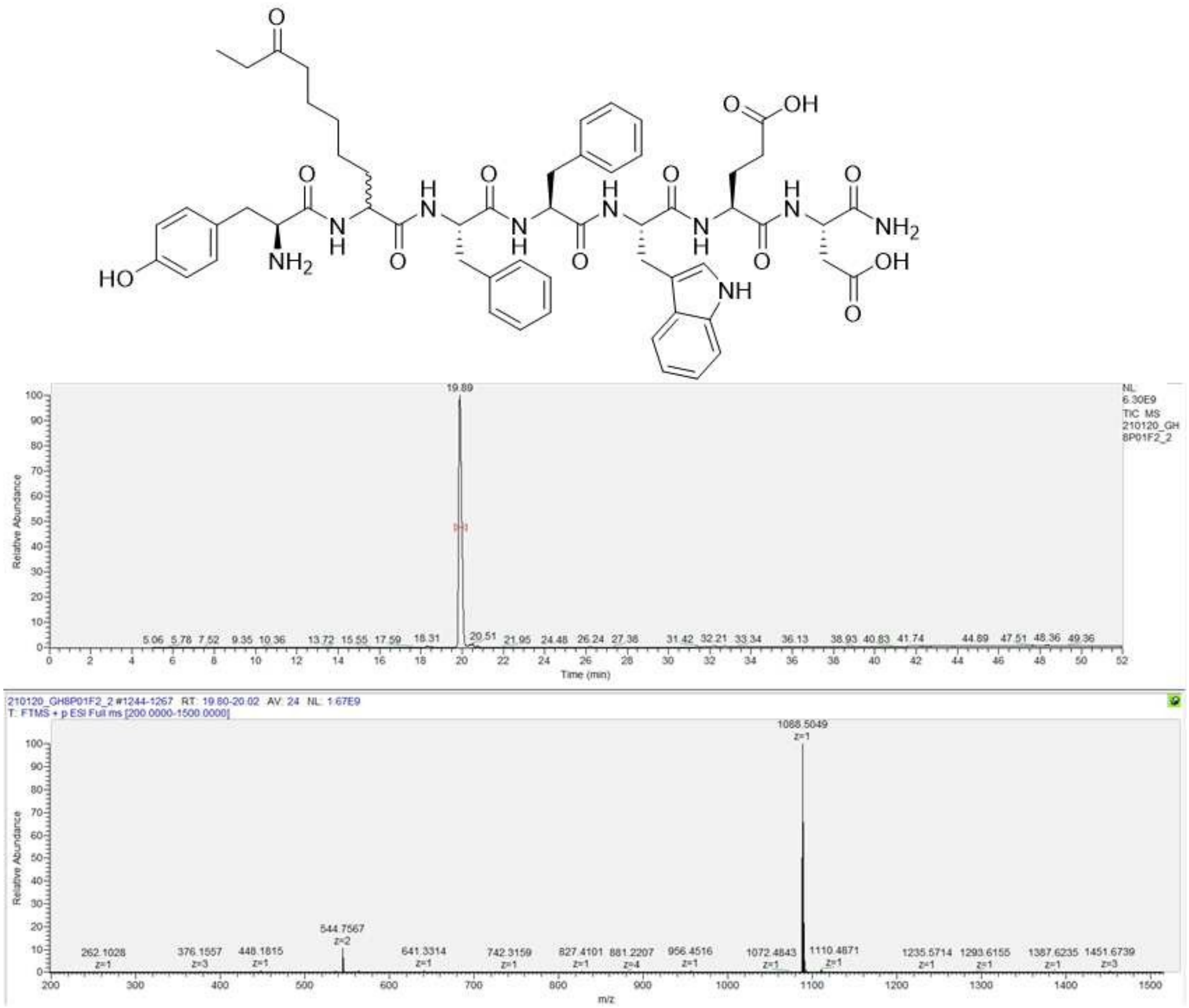


**Figure S5**: HPLC/MS of GH8P01F2. Calculated M+H$^+$: 1088.5088 Da.

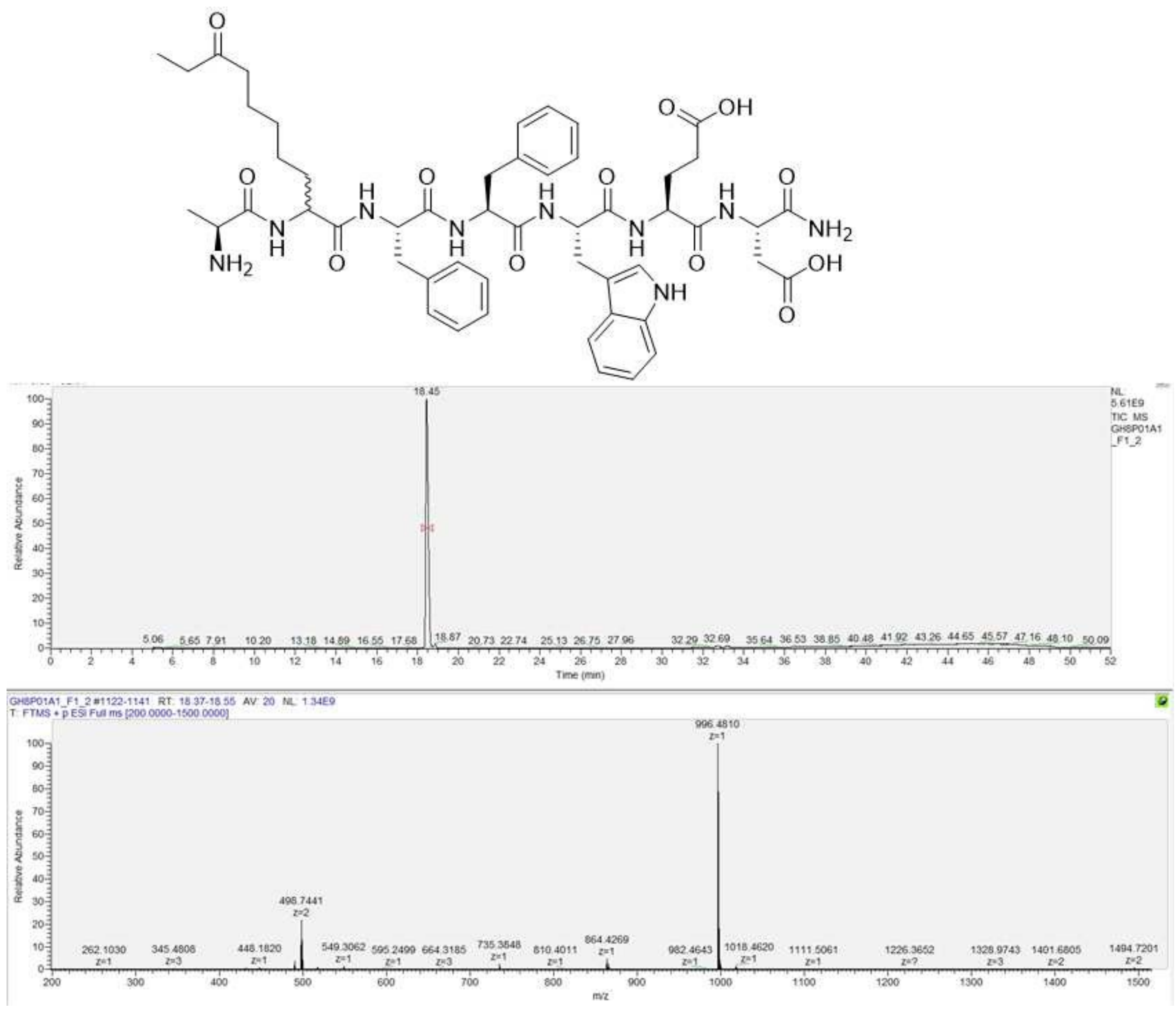


**Figure S6:** HPLC/MS of GH8P01A1F1. Calculated M+H$^+$: 996.4825 Da.

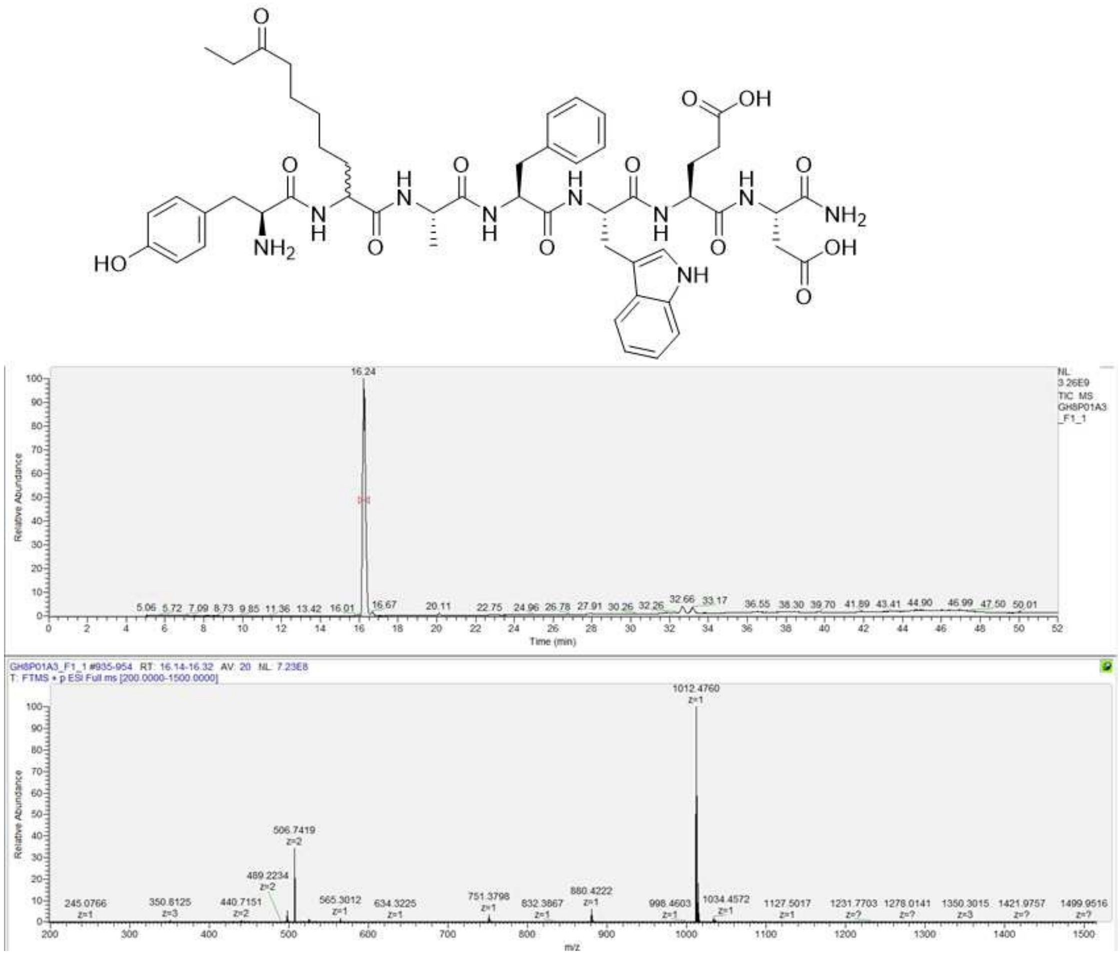

**Figure S7:** HPLC/MS of GH8P01A3F1. Calculated M+H$^+$: 1012.4775 Da.

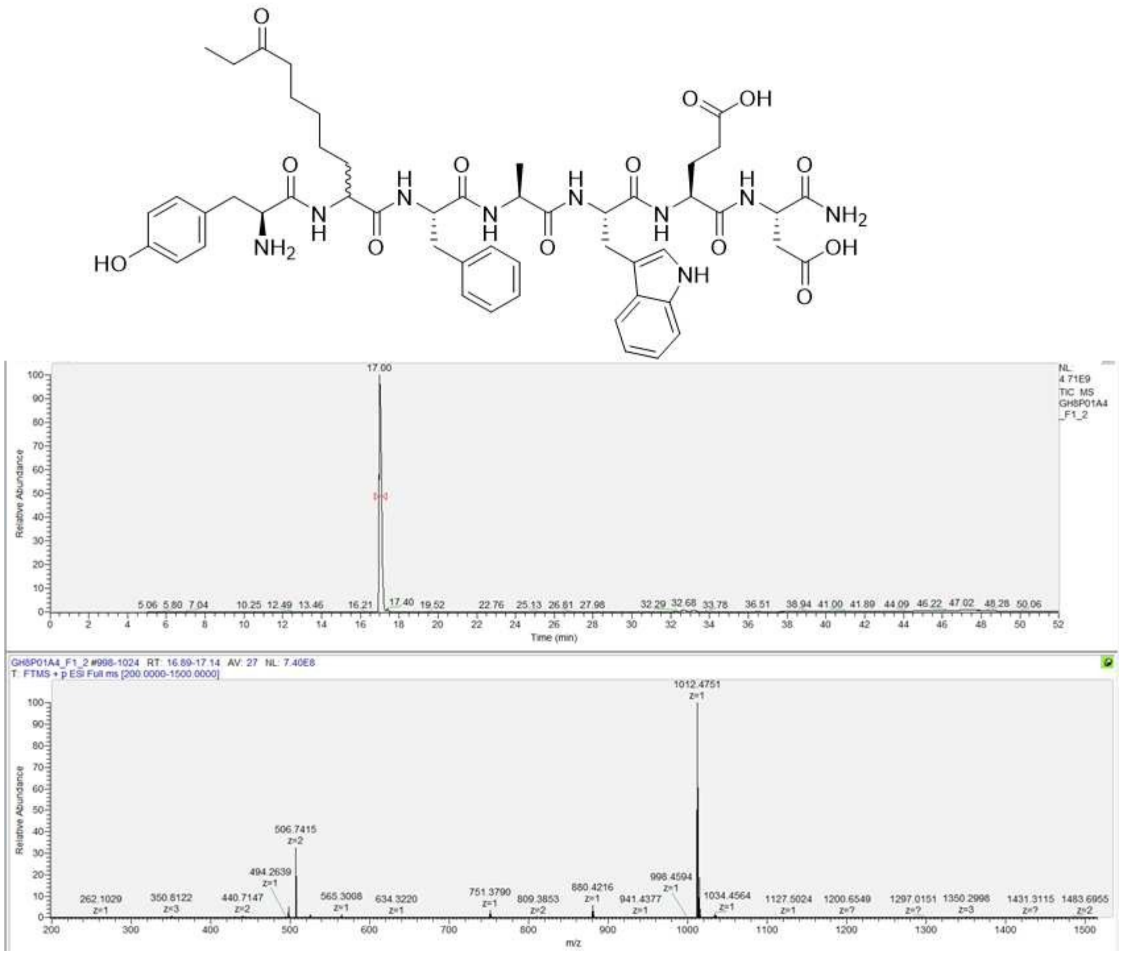


**Figure S8:** HPLC/MS of GH8P01A4F1. Calculated M+H$^+$: 1012.4775 Da.

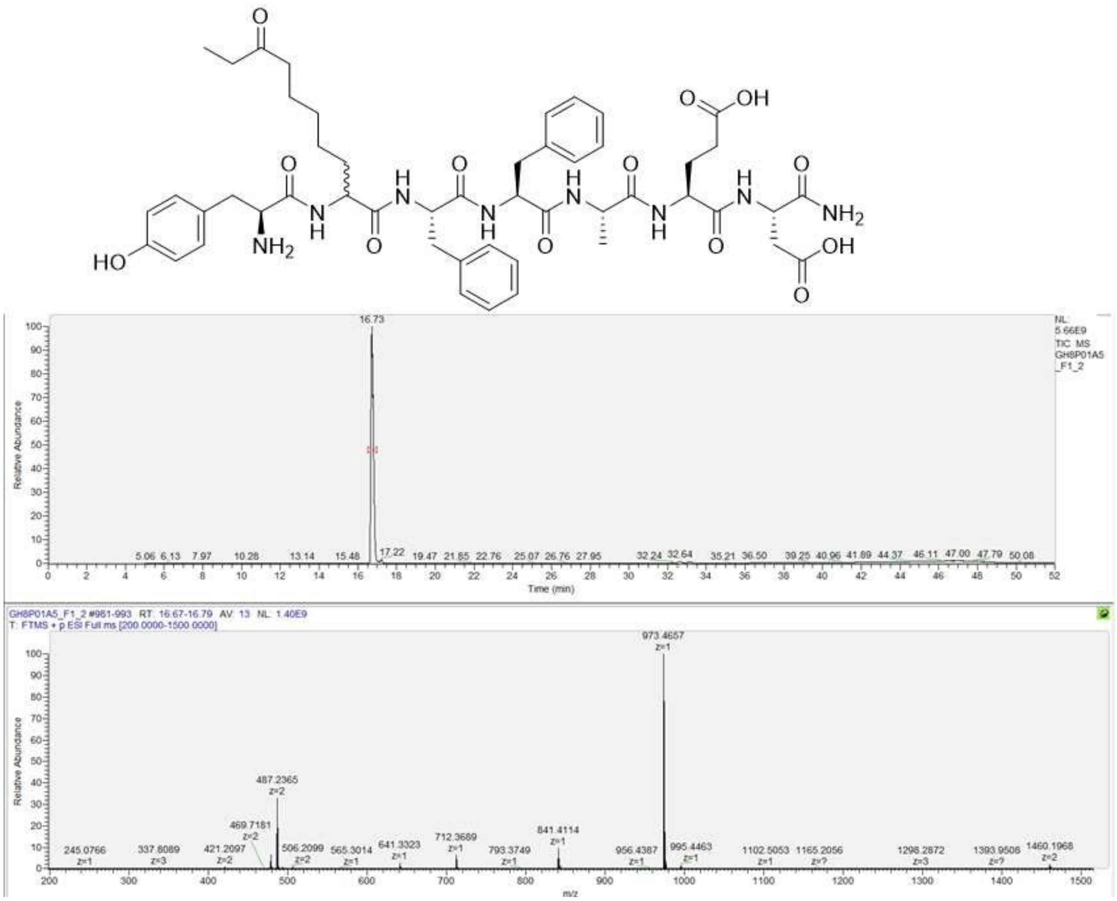

**Figure S9:** HPLC/MS of GH8P01A5F1. Calculated M+H$^+$: 973.4666 Da.

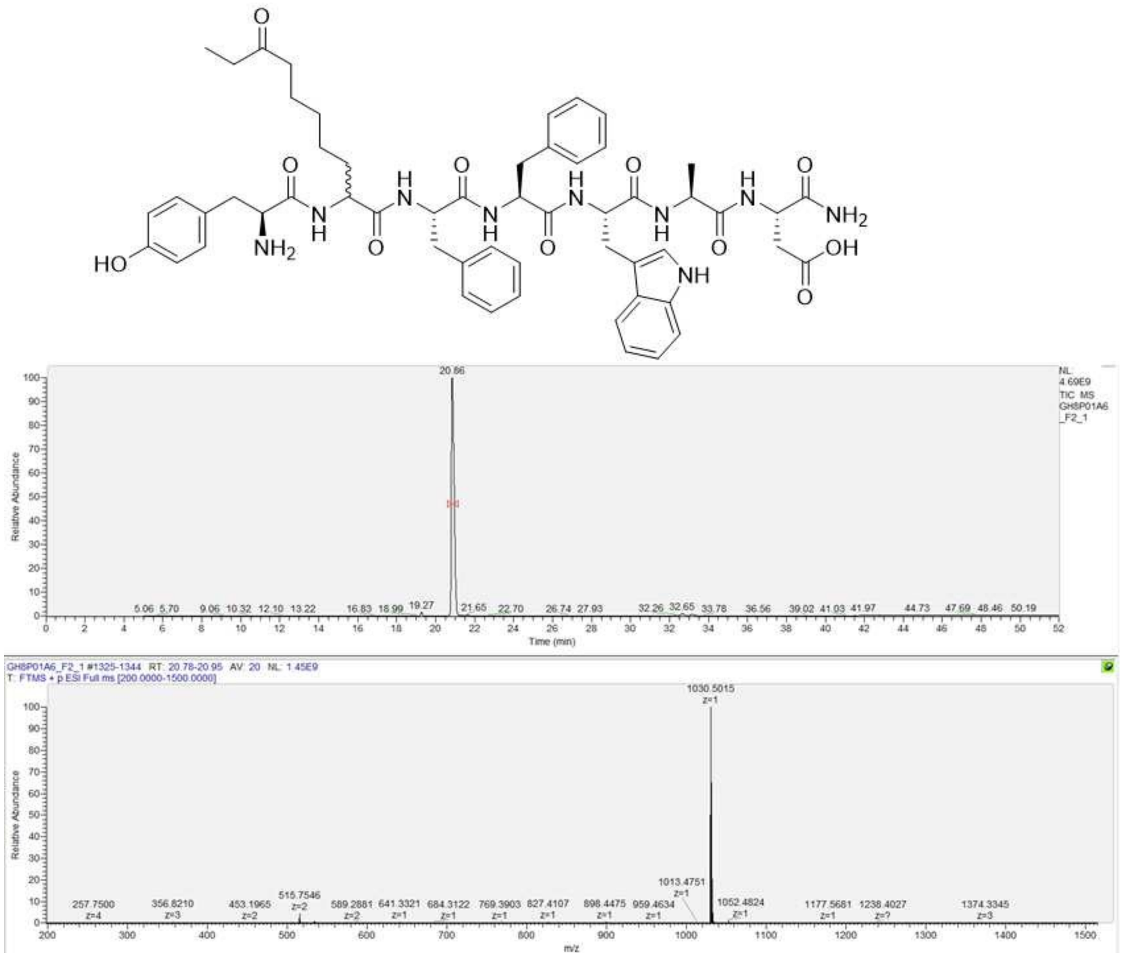


**Figure S10:** HPLC/MS of GH8P01A6F1. Calculated M+H$^+$: 1030.5033 Da.

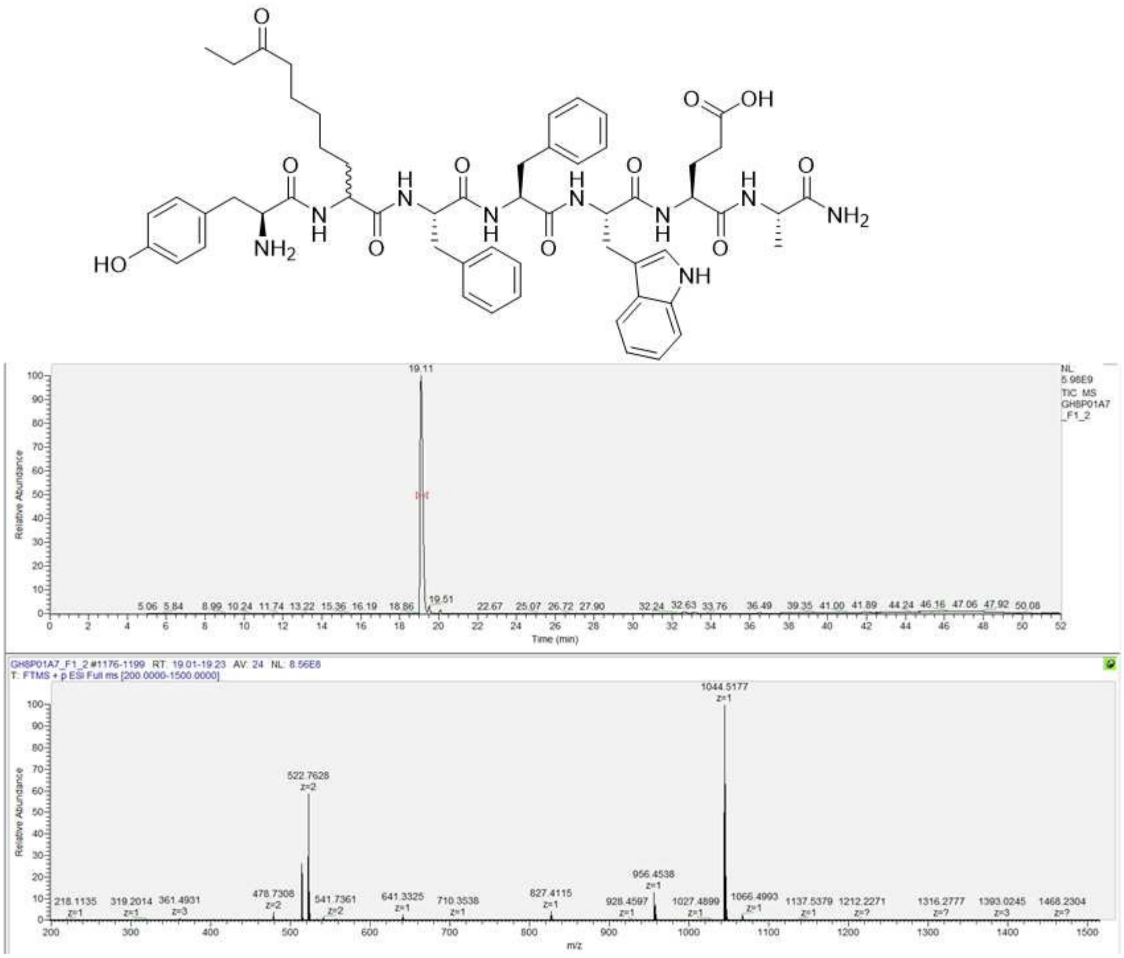


**Figure S11:** HPLC/MS of GH8P01A7F1. Calculated M+H$^{+}$: 1044.5189 Da.

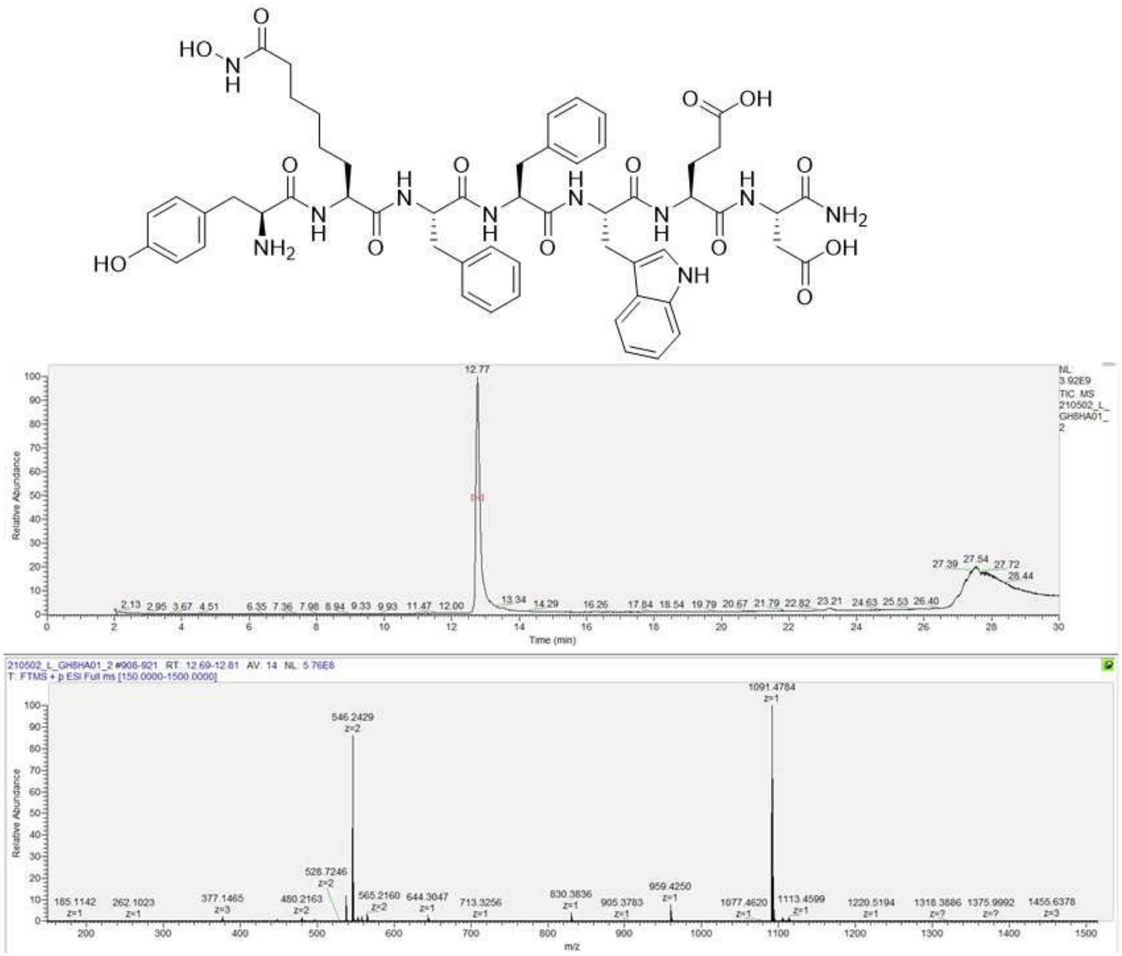


**Figure S12:** HPLC/MS of GH8HA01. Calculated M+H$^+$: 1091.4833 Da.

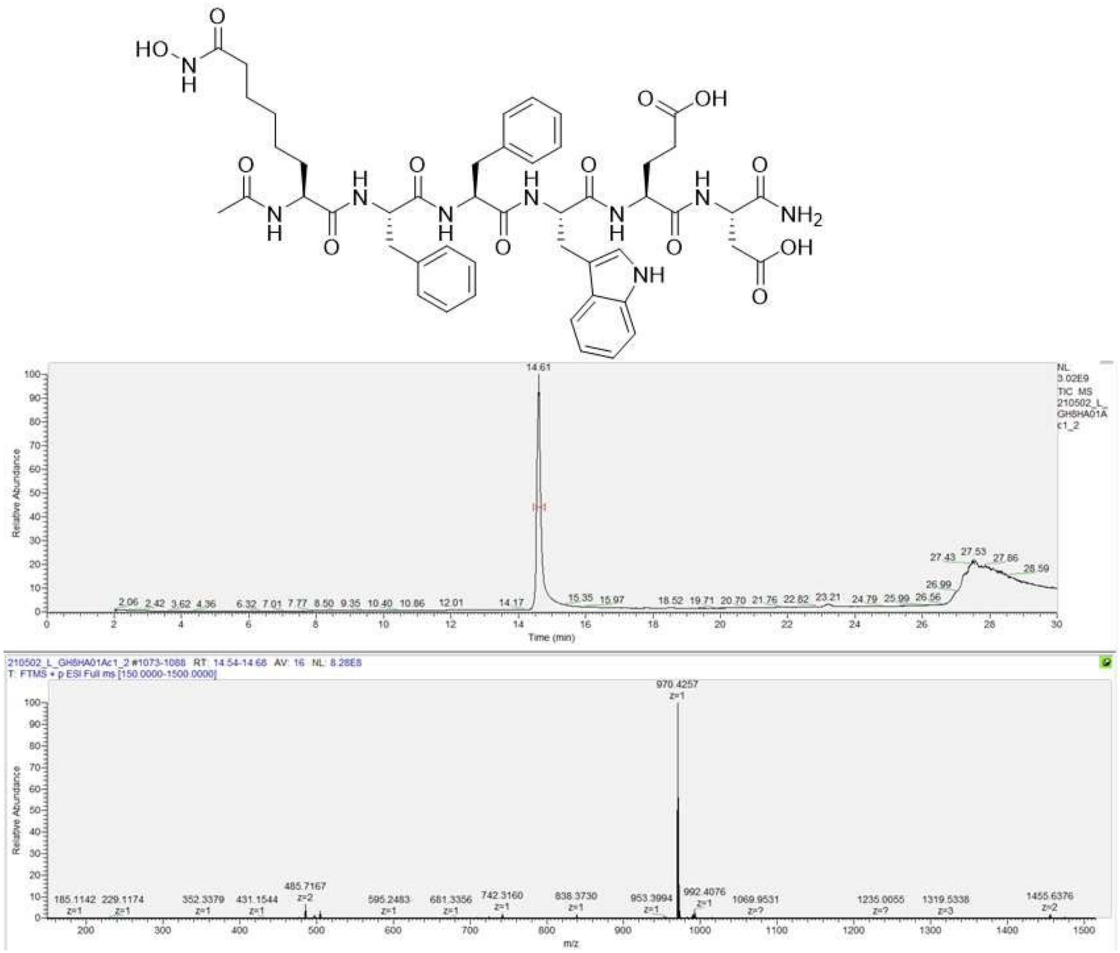


**Figure S13:** HPLC/MS of GH8HA01Ac1. Calculated M+H$^+$: 970.4305 Da.

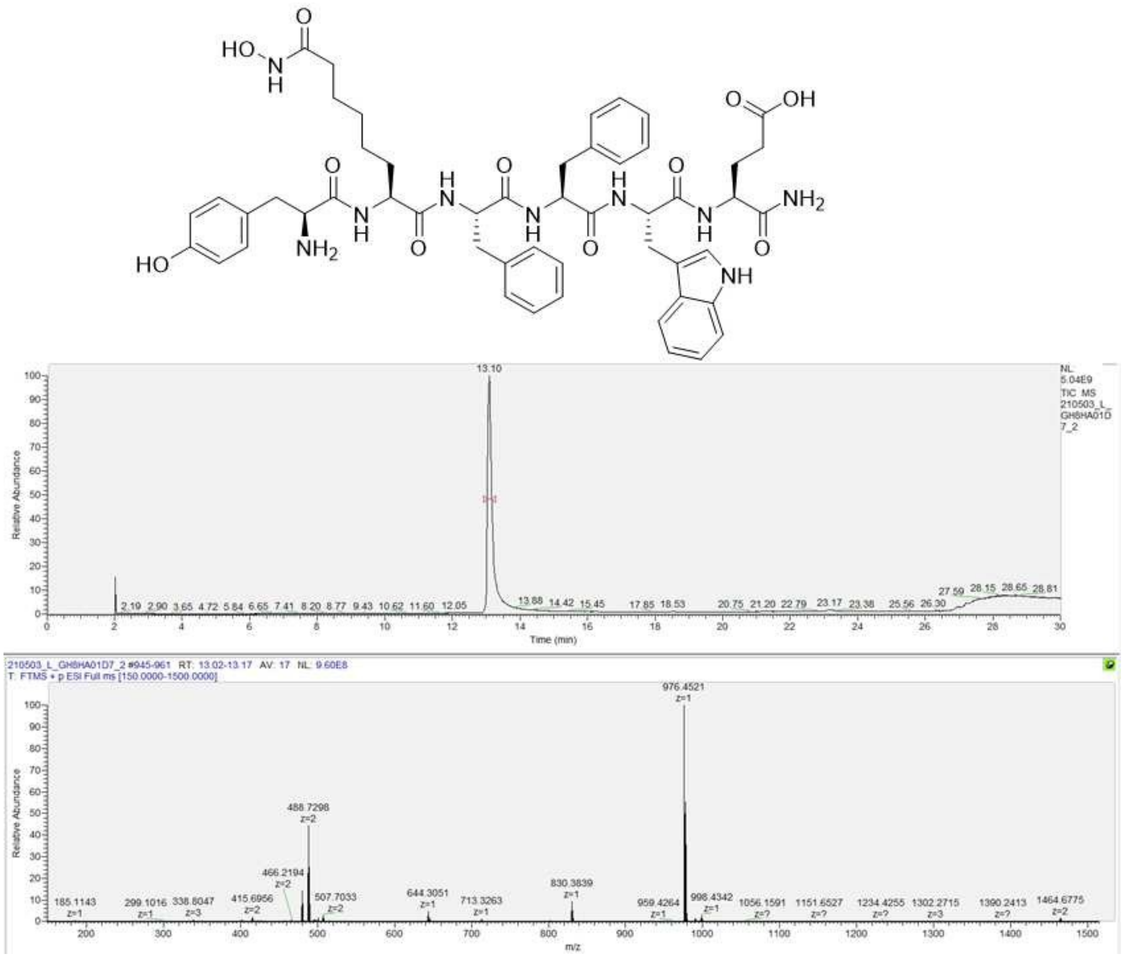

**Figure S14:** HPLC/MS of GH8HA01D7. Calculated M+H$^+$: 976.4563 Da.

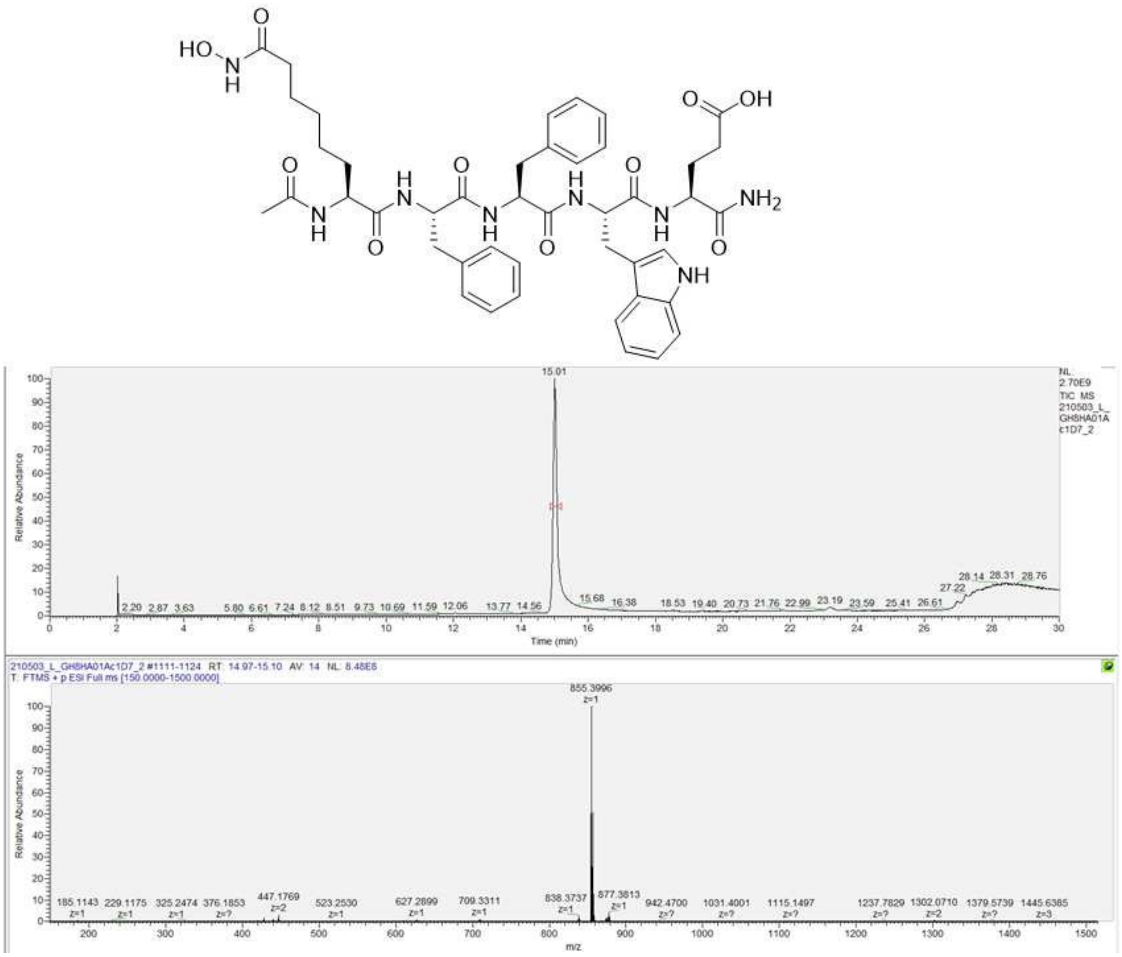


**Figure S15:** HPLC/MS of GH8HA01Ac1D7. Calculated M+H$^+$: 855.4036 Da.

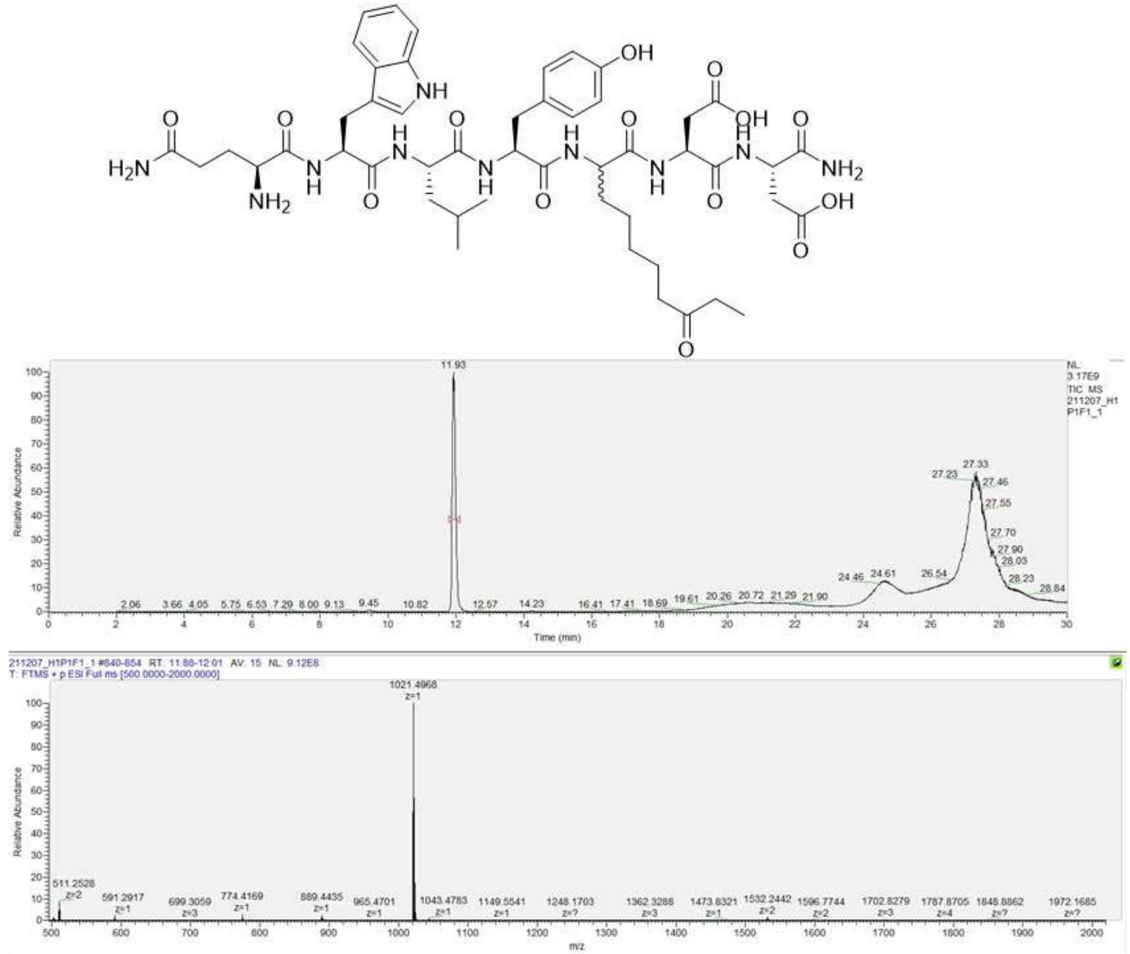


**Figure S16:** HPLC/MS of H1P01F1. Calculated $M+H^+$: 1021.4989 Da.

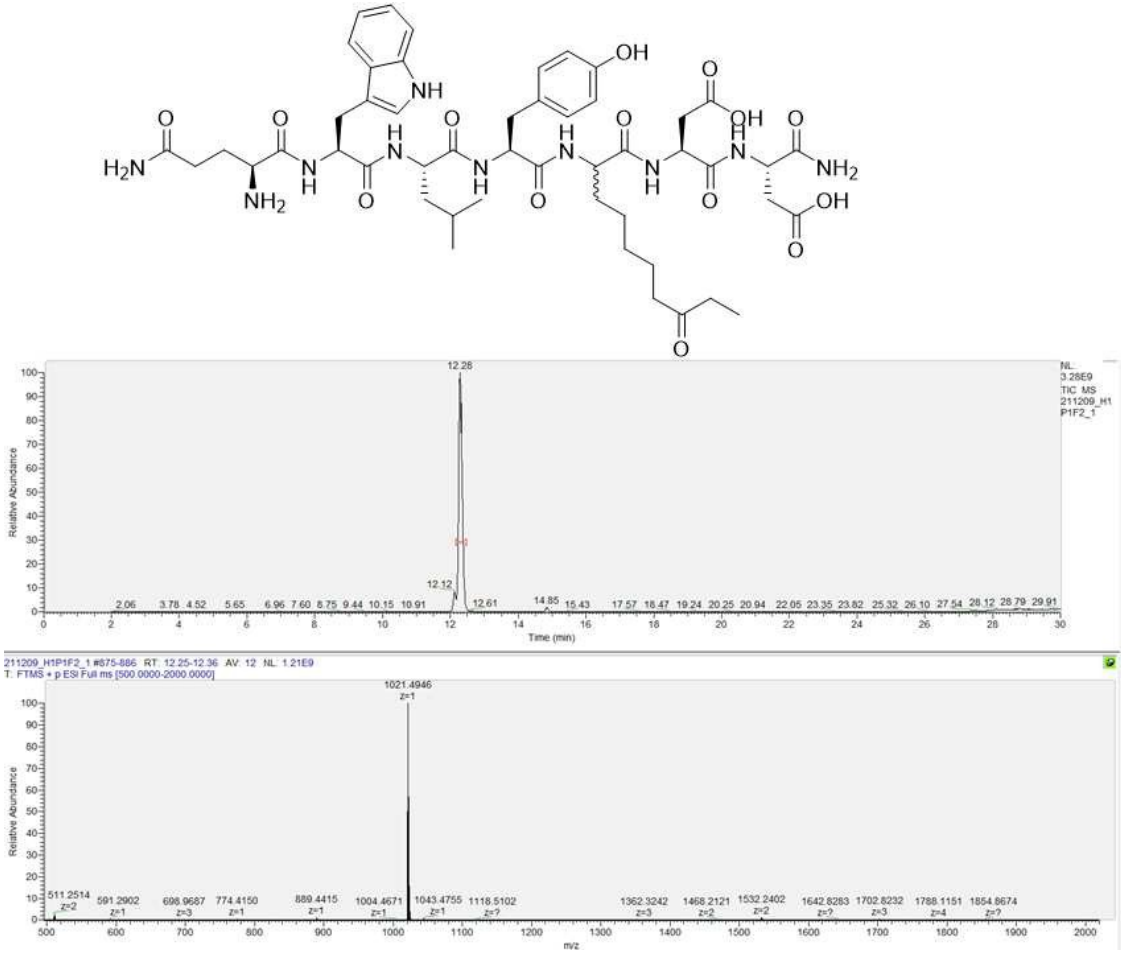


**Figure S17:** HPLC/MS of H1P01F2. Calculated M+H$^+$: 1021.4989 Da.

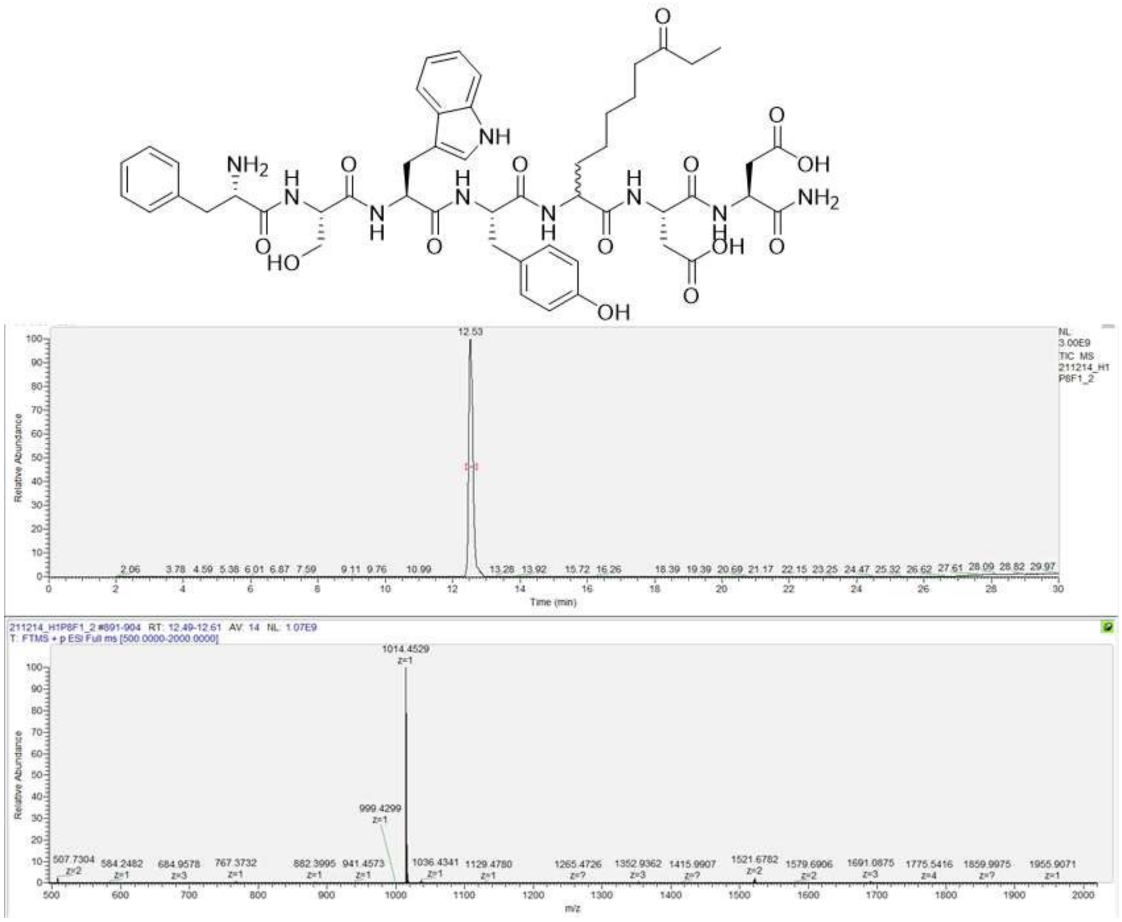


**Figure S18:** HPLC/MS of H1P08F1. Calculated M+H$^+$: 1014.4597 Da.

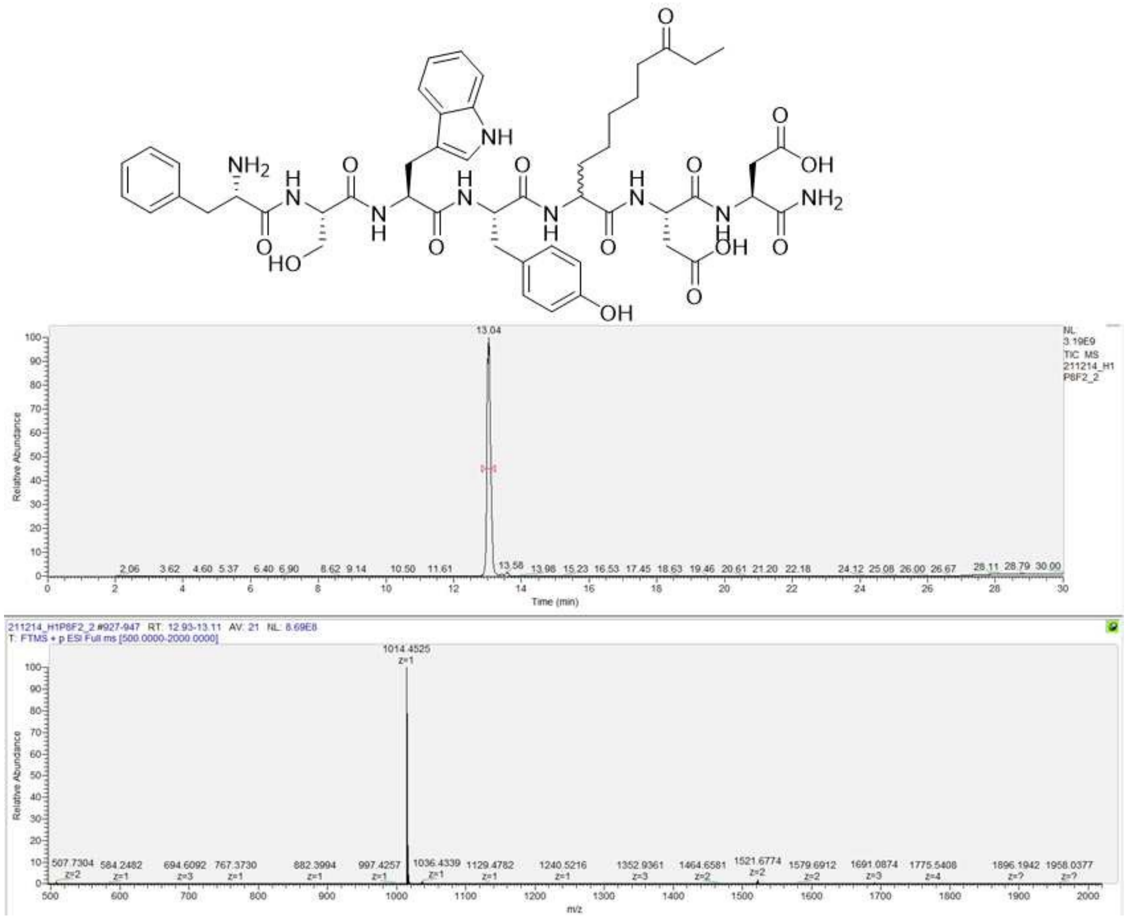


**Figure S19:** HPLC/MS of H1P08F2. Calculated $M+H^+$: 1014.4597 Da.
